# History of divergence and gene flow shaping geographic variation in Andean warblers (*Myioborus*)

**DOI:** 10.1101/2025.08.12.667688

**Authors:** Laura N. Céspedes Arias, Andrés M. Cuervo, Carlos Daniel Cadena, Elisa Bonaccorso, Christopher Witt, Irby Lovette, Leonardo Campagna

## Abstract

Studying how genetic variation is structured across space, and how it relates to divergence in phenotypic traits relevant to reproductive isolation, is important for our overall understanding of the speciation process. We used reduced-representation genomic data (ddRAD-seq) to examine patterns of genetic variation across the full distribution of an Andean warbler species complex (*Myioborus ornatus–melanocephalus)*, which includes a known hybrid zone between two strikingly different plumage forms. Genetic structure largely reflects geographic variation in head plumage, some of which corresponds to major topographic barriers in the Andes. We also found evidence of isolation by distance shaping genetic patterns across the group’s broad latitudinal range. We found that *chrysops* and *bairdi*, two taxa with marked plumage differences that have a known hybrid zone, were characterized by low overall genetic divergence. Based on our cline analyses of both plumage and genomic hybrid indices, this hybrid zone extends for approximately 250 km, where advanced generation hybrids are likely most common. We also identified a slight difference in the centers of the plumage and genomic clines, potentially suggesting the asymmetric introgression of *chrysops*-like plumage traits. By studying genetic variation in a phenotypically complex group distributed across a topographically complex area, which includes a hybrid zone, we were able to show how both geographic features and potentially sexually selected plumage traits may play a role in species formation in tropical mountains

## Introduction

Geographic variation in traits relevant to reproductive isolation is often viewed as a snapshot of the early stages of population divergence (Mayr, 1942; Price, 2008), making it a focus for understanding the process of speciation. To study this process, it is useful to evaluate how phenotypic variation in such traits is related to the underlying population genetic structure (Edwards et al., 2016; Zamudio et al., 2016) A common expectation is that phenotypic and genetic divergence are positively correlated: in the absence of strong divergent selection, extended periods of isolation are typically required for phenotypic differentiation to evolve (Nosil, 2008; Price, 2008; Winger & Bates, 2015). It follows that range disjunctions associated with topographic and ecological barriers likely to have restricted historical gene flow are expected to be accompanied by genetic and phenotypic discontinuities (Edwards et al., 2016). However, geographic variation in phenotypic traits is not always concordant with genetic structure. This discordance could be driven, for example, by strong selection on some traits across environmental gradients over short time scales (Walsh et al., 2017), or complex histories of selection and gene flow leading to phenotypic convergence (Márquez et al., 2020). A better understanding of how species diversify across space requires studying such histories of isolation, and how these relate to divergence in traits, particularly those that might be important for mate choice and, consequently, reproductive isolation.

Patterns of plumage differentiation—thought to be important for reproductive isolation in birds (Price, 2008; Uy et al., 2018)— among allopatric populations within species are pervasive among Andean birds (Graves, 1988; Remsen, 1984). Although geographic variation in plumage and genetic structure in Andean birds is often associated with current topographic and ecological barriers (Gutiérrez-Pinto et al., 2012; Prieto-Torres et al., 2018), many of these barriers tend to be permeable and unstable. Dispersal over such barriers could potentially lead to introgressive hybridization that connecting otherwise diverging populations (Winger, 2017). Additionally, differentiated populations can come into renewed contact when the barriers themselves change over time. In the particular case of the Andes, climatic fluctuations (Ramírez-Barahona & Eguiarte, 2013) and, potentially, volcanic activity (Sanín et al., 2024) have promoted cycles of fragmentation and reconnection of elevational vegetation belts, creating opportunities for contact among periodically isolated populations (Vuilleumier, 1969). Examining these complex histories of divergence and gene flow in geographically variable and widely distributed Andean taxa contributes to a more comprehensive understanding of how bird species are formed and maintained in this biodiversity hotspot (Rahbek et al., 2019; Sonne et al., 2022).

Here, we used a reduced representation genomic data set from an Andean bird species complex of *Myioborus* warblers (Parulidae) characterized by striking geographic variation in head color patterns to gain insights into differentiation in geographically isolated populations and the processes that occur after differentiated populations come into secondary contact. This young (<1 my; Cuervo, 2013) species complex inhabits the high Andes from Bolivia to Venezuela, and is characterized by differences in head color patterns despite shallow genetic differentiation in mitochondrial DNA (Céspedes-Arias et al., 2021; Pérez-Emán, 2005). Seven plumage groups have been described as subspecies belonging to two species, *Myioborus ornatus* and *M. melanocephalus*, distributed mostly allopatrically along different sections of the Andes (Cuervo & Céspedes Arias, 2023; Zimmer, 1949). According to this classification scheme, *M. ornatus* includes subspecies *ornatus* and *chrysops*, and *M. melanocephalus* includes subspecies *bairdi, griseonuchus, malaris, melanocephalus* and *bolivianus*. Recent taxonomic revisions, however, suggest that a four-species classification scheme might better capture what is known about genetic and plumage variation in this complex (Cuervo & Céspedes Arias, 2023). Regardless of the species delimitation scheme, analyses based on the ND2 mitochondrial gene have revealed extensive haplotype sharing among groups with conspicuous plumage differences, potentially suggesting a history of rapid plumage evolution or repeated mitochondrial introgression (Céspedes-Arias et al., 2021).

Our previous work in regions where the ranges of *chrysops* and *bairdi* abut, on the border between Colombia and Ecuador, revealed that these two taxa hybridize widely across more than 200 km where individuals with a variety of intermediate head color patterns occur (Céspedes-Arias et al., 2021). Although hybridization there has been extensive, the existence of a sharp plumage cline, limited to a relatively narrow zone within these taxa’s broader ranges (20‒30%), suggests the action of some form of selection maintaining their differentiation (Barton & Hewitt, 1985; Brelsford & Irwin, 2009; Harrison, 1986). Here, we leveraged this previous systematic collection and detailed plumage characterization of the *chrysops-bairdi* hybrid zone (Céspedes-Arias et al., 2021) to gain a deeper understanding of hybridization dynamics using genomic data.

We examined patterns of genomic variation within this warbler species complex at two different spatial scales: the entire distribution of the species complex, which corresponds to a broad latitudinal band in the Andes (approximately 3800 km in geographic distance), and the hybrid zone existing between two markedly different plumage groups as outlined above. By focusing on these two different scales, we can address questions related to multiple processes relevant to the formation and maintenance of species. First, leveraging a broad geographic scale sampling of individuals representing all plumage forms in the complex, we asked: (1) How is genetic structure related to geographic distance, topography, and plumage variation along the Andes? At a finer spatial scale, in the hybrid zone between two plumage forms, we asked: (2.1) Does the geographic pattern of introgression mirror plumage variation across the hybrid zone? And (2.2) is there evidence for non-neutral processes such as selection against hybrids based on patterns of genomic variation? Our genome-wide data coupled with dense geographic sampling substantially improves our ability to reconstruct and understand the diversification history of these colorful montane warblers, which was previously limited to studies using only a single mitochondrial gene.

## Methods

### Study species and sampling

We used tissue samples (*n*=227, Figure 1) collected across the distribution of the *M. ornatus-melanocephalus* complex, by us (Céspedes-Arias et al., 2021), or previously deposited in museum collections (Supplementary Table 1). Samples encompassed all (seven) currently taxonomically recognized plumage forms and a denser sampling in the *bairdi-chrysops* hybrid zone area, along the border between Colombia and Ecuador (Figure 1A). We also obtained data for eight individuals as outgroups corresponding to three other species in *Myioborus*: *M. pictus* (n = 3), *M. miniatus* (n = 3) and *M. brunniceps* (n = 2) which vary in their phylogenetic and geographic proximity to the *M. ornatus-melanocephalus* complex (Lovette et al., 2010; Pérez-Emán, 2005).

**Figure 1:**
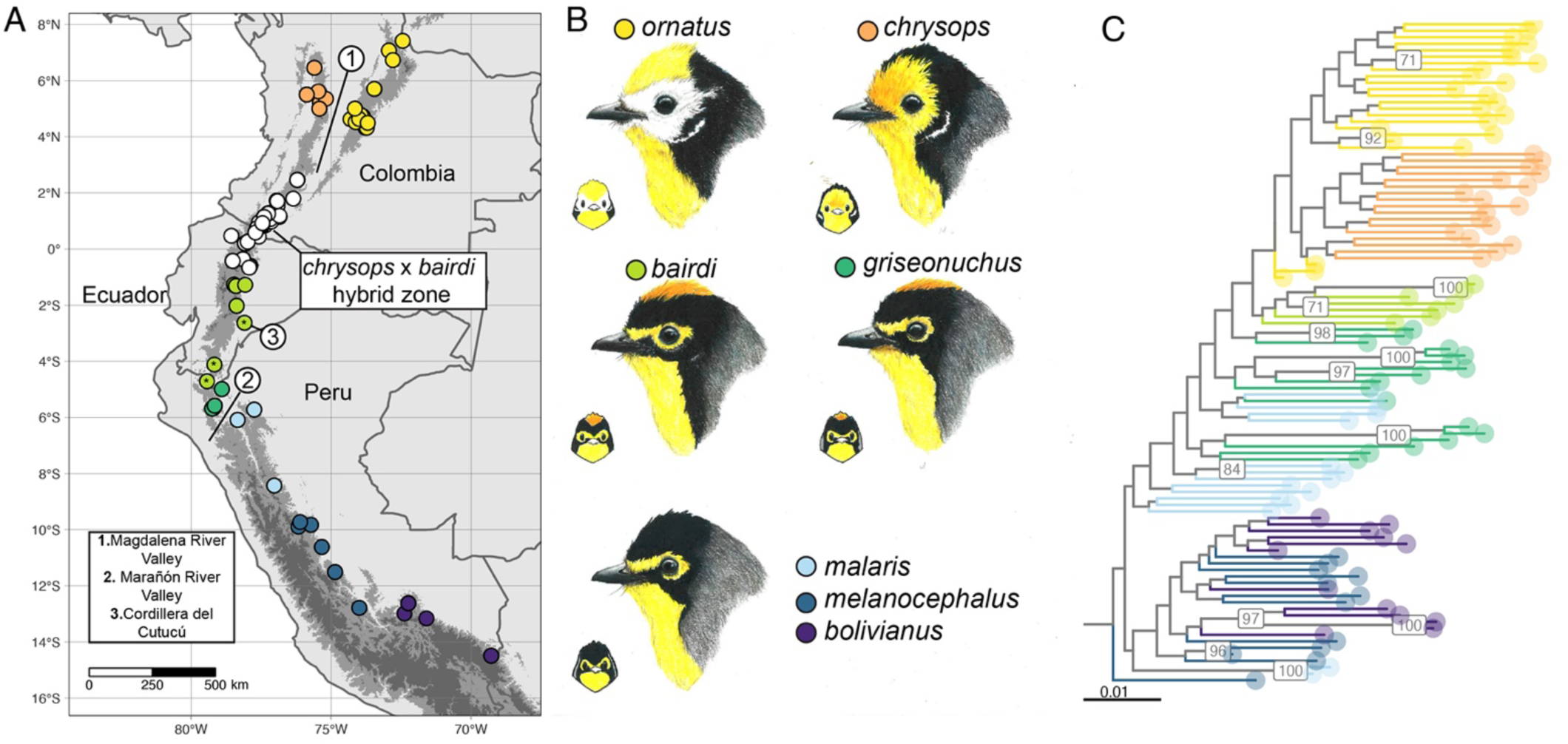
Sampling localities of specimens included in the study **(A)**, which encompass all major plumage groups **(B)**. The color of the dots corresponds to study taxa. The white dots correspond to areas around the *bairdi-chrysops* hybrid zone as identified based on plumage patterns (Céspedes-Arias et al., 2021). The subspecies *malaris, melanocephalus*, and *bolivianus* are here represented by one illustration. In the map, elevation is denoted by the gray scale in the background, with the elevation in which these *Myioborus* occur shown in an intermediate shade of gray. The localities marked with an asterisk correspond to those in Loja and outlining ridges in Morona-Santiago that are discussed in the main text, and numbers mark important topographic features. **(C)** Maximum-likelihood phylogeny estimated with our dataset of concatenated ddRAD loci, showing how overall relationships somewhat reflect geography (outgroups are cropped out of the figure here). The branches are colored by subspecies as in the map, with only bootstrap values over 70 shown (hybrid zone samples not included in the analysis). Illustrations: Andrés Montes Rojas.

The more conspicous differences among plumage groups in the *ornatus-melanocephalus* complex correspond to the presence/absence and extent of colored plumage patches in the head, as illustrated in Figure 1B. We note that there are different degrees of plumage differentiation and diagnosability based on our own assessments (Cuervo & Céspedes Arias, 2023). For example, the subspecies *malaris, bolivianus* and *melanocephalus* are very similar in plumage, and some of the described diagnostic traits do not seem to vary in a way that is discrete and corresponds with these groups (Cuervo & Céspedes Arias, 2023).

We also note that the geographic limits of some groups are also not clearly defined, specifically of the *bairdi* and *griseonuchus* groups which have only subtle differences in in the amount of yellow vs black on the sides of the face, as well as the presence and extent of a black nuchal band behind the rufous crown (Chapman, 1927; Cuervo & Céspedes Arias, 2023; Zimmer, 1949). Although birds from northern Peru (Dept. of Cajamarca) and central Ecuador are clearly distinct (Supplementary Figure 1), it is unclear whether a phenotypic transition is discrete or clinal across intervening southern Ecuador and there is a lack of sampling across this area to evaluate these possibilities. However, to inform our discussion on patterns of genetic variation, we inspected most of the available specimens collected in this area, either directly or through photographs, to describe variation in these two plumage traits (Supplementary Figure 1).

### Laboratory procedures

#### DNA extraction and library preparation

We extracted genomic DNA from pectoral muscle (n=232) and blood samples (n=3) employing a modified phenol-chlorophorm protocol (Sambrook, 1987), using phase-lock tubes. Each DNA extraction was subsequently purified using homemade Sera-Mag Magnetic Speed-beads for magnetic nucleic acid extraction (MagNA). To obtain reduced representation genomic data, we used double-digest restriction site-associated DNA sequencing (ddRAD) following Peterson et al. (2012), and using modifications from Thrasher et al. (2018). The genomic libraries were then sequenced on one HiSeq2500 Illumina lane (100 bp SE) by the Genomics Core Facility of Cornell University.

### Bioinformatics pipeline

#### Quality assessment, filtering and demultiplexing

We assessed sequence quality using FastQC (Andrews, 2010). We used the *process_radtags* function implemented in STACKS 2.66 (Rochette et al., 2019) to demultiplex the reads and apply quality filters. Specifically, we filtered out all reads with uncalled bases (*--clean*), those with low phred quality scores *(--quality*, *--score-limit* 20) and those that did not pass the Illumina chastity/purity filter (*--filter-illumina*). After demultiplexing, we checked the number of reads available per individual, and only kept those samples with 200,000 or more reads (Supplementary Figure 2). Using this threshold, we discarded 11 individuals, mostly corresponding to the *ornatus* group and tissues collected around ten years prior to this study (range of number of reads for these individuals = 960‒150,152). Despite discarding these samples, 23 high-quality samples for the *ornatus* group remained in downstream analyses. After excluding these individuals from downstream analyses, we kept a total of 242,204,894 reads corresponding to 224 individuals (three represented twice in the library).

#### Integrated assembly and SNP calling

After demultiplexing, we assembled the RAD reads *de-novo* using functions implemented in STACKS 2.66 (Rochette et al., 2019), and later integrated position information for the loci in the catalog from a reference genome (Paris et al., 2017; Rivera-Colón & Catchen, 2022). This integrated approach performs better than directly mapping reads onto a reference genome in some scenarios, for example, if the reference genome is relatively distant to the study species as is our case (Paris et al., 2017). We used the chromosome-level assembly of the *Setophaga coronata* genome (Baiz et al., 2021) as reference, the highest quality available genome within Parulidae. The time to the most recent common ancestor between *Setophaga* and *Myioborus* is estimated between 5‒7 mya (Barker et al., 2015).

#### Optimization of de novo assembly parameters

Before assembling the reads, we conducted an optimization of the parameters *M* and *n* used by *denovo_map.pl* (Rochette et al., 2019), for three different assemblies: one including representatives from the outgroups, one that only included individuals within the *ornatus-melanocephalus* complex, and one including only samples relevant for the hybrid zone analyses (*chrysops*, individuals in the hybrid zone, *bairdi,* and *griseonuchus*, Figure 1) (see Supplementary Material and Supplementary Table 2 for details).

#### Full de novo assembly run

We then ran *denovo_map.pl* for the entire dataset with the chosen parameter values, using again the three population maps (n = 224 full assembly, n = 216 ingroup only assembly, and n = 123 for the hybrid zone assembly). For the population maps corresponding to the three different assemblies, we classified individuals into one of the following groups: *ornatus*, *chrysops*, “hybrid zone”, *bairdi, griseonuchus, malaris, melanocephalus*, and *bolivianus* (populations_2 column in Supplementary Table 1), plus *pictus, miniatus*, and *brunniceps* for the assembly including the outgroups. For the population map purposes the hybrid zone group corresponds to a geographic group encompassing populations in and near the hybrid zone, irrespective of the plumage phenotype of the specimens (see the map in Figure 1).

After the assemblies were completed, we assessed the per sample coverage and percentage of reads incorporated (Rivera-Colón & Catchen, 2022). The assembly corresponding to only individuals within the *melanocephalus-ornatus* complex resulted in a catalog of 91,650 loci, with a mean effective coverage per sample of 59.1x (SD = 13.2x, min = 18.3x, max = 97.4). The “hybrid zone” assembly resulted in a catalog of 70,281 loci, with a mean effective coverage per sample of 61.6x (SD =12.1x, min = 22.5x, max = 99.4x). The assembly including outgroups resulted in a catalog of 97,533 loci, with a mean effective coverage per sample of 59.3x (SD = 13.2x, min = 18.1x, max = 98.5x). Four individuals with a mean effective coverage lower than 20x in assemblies in which they were included initially were excluded from downstream analyses (see Supplementary Table 1; Supplementary Figure 3).

#### Mapping to the reference genome

After obtaining the *de novo* catalog, we mapped loci to the reference genome by first indexing the *S. coronata* genome using *bwa index* (Li & Durbin, 2009). Then we used *bwa mem* to align the *de novo* catalog to the reference genome. Finally, we used the *stacks-integrate-alignments* script to generate a catalog with position information (Rivera-Colón & Catchen, 2022; Rochette et al., 2019). After filtering based on mapping quality and alignment coverage (using default values), 60,523 loci (66% of the total number of loci in *de novo* catalog) were retained for the ingroup assembly, 46,036 (66% of the total number of loci in *de novo* catalog) were retained for the “hybrid zone” assembly, and 65,537 (67% of the total number of loci in *de novo* catalog) for the full assembly.

#### SNP datasets

We obtained different SNP datasets to be used in downstream analyses using the *populations* program from STACKS 2.66 (Rochette et al., 2019). For some analyses, as specified below, we did further filtering of SNPs using *vcftools* (Danecek et al., 2011). For the ingroup assembly, we obtained two different SNP datasets: one including one SNP per locus, and one that included all variant and non-variant sites per locus (i.e., full haplotypes only for fineRADStructure, see below). In both cases we applied the following filters: *--min-samples-overall* 0.8 (locus needs to be present in 80% of all individuals to be retained), *--min-mac* 5 (minimum allele count of 5), and *--max-obs-het* 0.7 (maximum observed heterozygosity of 0.7). For the full haplotype dataset, we used the same parameters and included the *--filter-haplotype-wise* flag. We decided to apply a minimum allele count (*--min-mac* 5) filter to exclude SNPs that may be artifacts of genotyping errors, but kept a relatively low value to avoid excluding all low frequency SNPs which also contain relevant information for population genetic analyses (Linck & Battey, 2019). Details on how the SNPs datasets from the three different assemblies were obtained are in the Supplementary information. In short, we used the same filtering parameters for the hybrid zone assembly and for the assembly including outgroups we used *--min-mac 2* and *--min-samples-per-pop* 0.5

Based on the assembly for the ingroup only, and after filtering, we obtained 5,054 loci, composed of 73,786 sites, including 22,132 variant sites for the full haplotype dataset. For the ingroup (one SNP per locus) dataset, we obtained the same number of loci and kept 4,800 variant sites.

### Analyses of diversification within the *ornatus-melanocephalus* complex

We described and visualized genetic structure in order to test its relationship with plumage and geography. To evaluate how genetic structure related to current taxonomic patterns of variation in head color patterns (see Figure 1), we also inferred subspecies-level phylogenies and admixture graphs. Additionally, we estimated the level of differentiation between plumage groups in terms of F_ST_. Lastly, we used a spatially explicit model to evaluate how genetic variation within the *ornatus-melanocephalus* complex is structured across their range.

#### Phylogenetic and admixture graph inference

With the goal of inferring the overall relationships among plumage groups, we employed two methods to build phylogenies and infer admixture graphs. For this set of analyses, we excluded all individuals from or adjacent to the hybrid zone (in white, Figure 1) and the *griseonuchus-bairdi* transition area (arrows, Figure 1), and used the assembly that includes outgroups (*M. pictus, M. miniatus, M. brunniceps*). For phylogenetic approaches, we included all outgroup individuals (total n=110), while for the admixture graph inference, we only included one outgroup (total n=105).

##### raxml-ng

We used RAxML-NG (Kozlov et al., 2019) to infer maximum-likelihood trees from concatenated ddRAD loci, including both variant and invariant sites. Because individual fragments are short (∼150 bp), we concatenated all loci into a supermatrix (Leaché & Oaks, 2017). A VCF including invariant sites was generated with *populations* using the parameters *-- vcf-all --min-mac* 2 *--min-populations* 10 *--min-samples-per-pop* 0.5, then converted to Nexus with *vcf2phylip*. (https://github.com/edgardomortiz/vcf2phylip). Then, we ran *raxml-ng* using this nexus alignment and the GTR + G model of nucleotide substitution with no partitions. To search for the tree with the highest likelihood, we ran 20 independent runs, 10 of them starting from the maximum parsimony tree and 10 from random trees. From these searches, the tree with the highest likelihood was selected among the 20 slightly different trees. We calculated branch support with 100 bootstrap replicates.

##### Snapper

To infer evolutionary relationships among plumage groups under a phylogenetic framework, we used *snapper* (Stoltz et al., 2021). *Snapper* infers a “species” tree using diffusion models and takes as input biallelic SNPs that are assumed to be unlinked, bypassing the necessity of building individual gene trees, similar to site-based coalescent methods (Bryant et al., 2012). We classified individuals into groups corresponding to the seven subspecies described in Figure 1. For this analysis, we generated a SNP dataset from the assembly including outgroups (*--min-mac* 2, *--max-obs-het* 0.7, --min-populations 10, *--min-samples-per-pop* 0.5) filtered to only keep biallelic SNPs using *vcftools* and then converted the resulting *vcf* files to nexus using the *vcf2phylip* script. Using the nexus alignments as input, we prepared the *.xml* files in Beauti 2.7.6 (Bouckaert et al., 2019). We ran four independent chains with a length of 15,000,000 steps, and a thinning interval of 1,500. We set a gamma prior for the birth rate (*λ*), and left default values for the remaining priors. We increased the weights of two operators (gamma mover and rate mixer) to 10. Stationarity was assessed using Tracer 1.7.1 (Rambaut et al., 2014), based on a visual exploration of trace plots and ESS values (ESS > 500 was obtained for all parameters.) After exploring trace plots, we set the burn-in at 10%.

##### TreeMix (OrientAGraph)

As an alternative way to describe the historic relationships among plumage groups, we used *OrientAGraph* (Molloy et al., 2021), an approach based on the *TreeMix* algorithm, which implements a graph-based model that uses genome-wide allele frequencies to infer population splits and admixture events (Pickrell & Pritchard, 2012). This method detects evidence of historical introgression between plumage groups and can better describe the history of populations when not fully bifurcating. For input, we applied the same SNP filtering as in the *snapper* analysis, adjusting only *--min-populations* to 8 and excluding SNPs with >30% missing data using *--max-missing* 0.7 in VCFtools. The filtered VCF was converted to TreeMix format with *vcf2treemix.py*. (https://github.com/CoBiG2/RAD_Tools/blob/master/vcf2treemix.py).

We ran *OrientAGraph* testing for a range of migration edges (*m* parameter), running 5 iterations per *m*, with *m* values ranging from 1 to 6. We used *M. miniatus* (n = 3) as outgroup to root the tree and incorporated a *bairdi-chrysops* known migration event using the -*cor_mig* and *- climb* flags, and a 0.5 migration weight. This was done to account for the high likelihood that there are signs of introgression even away from the hybrid zone. After running all iterations for this range of m values, we used *OptM* (Fitak, 2021) to evaluate the optimal value of *m* in each case, using an approximation based on linear regressions between *m* values and likelihood. We made a small modification to the *optm* function to be able to account for the known migration event. Based on this linear regression criteria, we chose the optimal *m* values and included those results here.

#### Genetic structure within the ornatus-melanocephalus complex

##### PCA

To summarize genetic variation in the *ornatus-melanocephalus* complex, we conducted a genomic PCA using functions implemented in *gdsmt* and *SNPrelate* (Zheng, 2013) for R. To conduct this PCA, we used only one SNP per locus, including only biallelic SNPs. To account for the effect of missing data on the PCA (Yi & Latch, 2022), we (1) visualized the amount of missing data in our PC plots, to evaluate the possibility that high missing data samples are clustered together and/or have values biased towards the origin, (2) ran an additional PCA only including SNPs with a missing rate < 0.1, and (3) ran an additional PCA using a different *vcf* as input with more strict filtering for missing data when running populations (*--min-samples-overall 0.9*). We also conducted a PCA for a subset of the individuals to describe in more detail genetic variation in the hybrid zone and adjacent areas (*chrysops*, hybrid zone individuals, *bairdi*, and *griseonuchus*).

##### fineRADStructure

As a complementary approach to visualize genetic structure within the *ornatus-melanocephalus* complex, we used *fineRADStructure*, which estimates a coancestry matrix and an associated distance tree based on full haplotype information from RADseq data (Malinsky et al., 2018). This method has particularly high resolution in inference of recent shared ancestry, and allows one to describe nested patterns of genetic structure (Malinsky et al., 2018). To run *finestructure* (part of the *fineRADStructure* pipeline), we used the output from the populations program containing both variant and invariant sites (i.e., full haplotypes). No population prior was specified for the run, which was set for 200,000 iterations with a burn-in of 100,000 (-x 100,000, -y 100,000), and a thinning interval of 1,000 (-z 1000).

##### Calculation of F_ST_

To estimate overall genetic differentiation among plumage groups, we calculated pairwise F_ST_ between subspecies (i.e., plumage and geographic groups). We excluded all individuals from the hybrid zone from this analysis. Given the uncertainty in the *bairdi-griseonuchus* limits (see Results), we also excluded individuals from the southern Ecuadorian provinces (as in the *snapper* analyses). We calculated the Weir and Cockerham estimator (Weir & Cockerham, 1984) using *vcftools* (Danecek et al., 2011) in 25,000 bp windows. To test for potential discordance between nuclear and mitochondrial divergence, we evaluated whether SNPs-based pairwise F_ST_ values were correlated with those estimated from the ND2 gene (Céspedes et al. 2021). To assess the correlation between these two matrices, we implemented a Mantel Test based on a Pearson’s correlation using the function *mantel* implemented in the R package *vegan* (Oksanen et al., 2007)

To test for isolation-by-distance (IBD), we also calculated pairwise windowed F_ST_ between localities as defined in Supplementary Table 1 and evaluated if these are correlated with the pairwise geographic distances. Geographic distances between localities were calculated using the function *rdist.earth* implemented in the R package *fields* (Nychka et al., 2025). We also evaluated the correlation between these matrices using a Mantel test based on a Pearson’s correlation as implemented in *vegan*.

##### Admixture

We employed admixture (Alexander et al., 2009) to estimate individual genetic ancestries using a maximum-likelihood approach at three different geographic scales: whole complex, hybrid zone and adjacent populations (*bairdi, chrysops*, hybrid zone, *griseonuchus*), and hybrid zone only (*bairdi*, *chrysops*, hybrid zone).

Prior to running admixture, we calculated relatedness between all individuals and then excluded individuals that appeared closely related. Specifically, we used the *--relatedness2* flag in *vcftools* to export a table of pairwise relatedness values using the KING kinship estimator (Manichaikul et al., 2010). We then identified pairs of individuals with a relatedness higher than 0.177 (i.e. likely first-degree relatives) and excluded one of them from admixture analyses. We then transformed the VCF files to *.bed* format by using plink2 (Chang et al., 2015). Additionally, we performed a SNP window linkage-disequilibrium pruning using plink2 *(--indep-pairwise*, followed by *--extract*), on windows of 50 SNPs and using a 0.3 threshold. This was done for both scales of analyses, and then admixture was run on the thinned datasets.

For all admixture runs, we included the *–cv* flag, to perform the cross-validation procedure to choose the optimal *k* (i.e. number of source populations) value. For the whole complex analyses, we ran admixture for values of *k*{1…10}, and for the other two scales, we used values of *k*{1…6}. For each value of *k*, we ran 10 iterations starting with a randomly generated seed. We analyzed variation across runs using *pong* (Behr et al., 2016) and generated figures using the *mapmixture* (Jenkins, 2024) package for R.

##### FEEMS

To more directly evaluate how genetic variation is shaped across the Andes, we used a spatially explicit model based approach: *FEEMS* (Marcus et al., 2021) (https://github.com/VivaswatS/feems). This approach is based on a model of IBD and builds continuous migration surfaces based on a geographic grid, visualizing areas of inferred low and high gene flow (Marcus et al., 2021). To run FEEMS, we prepared four input files: (1) *plink* file with genotype data for all individuals in the *ornatus-melanocephalus* complex (same set as for whole-complex admixture analyses), (2) text file with geographic coordinates for all specimens with genotype data (Supplementary Table 1), (3) a global grid with triangular cells of approximately 7,774 km^2^, obtained using the package *dggridR* for R (res=8; Barnes et al. 2024), and (4) outer extent boundary for the grid, which was generated as a polygon that roughly delineates the montane areas occupied by *Myioborus* and encompasses all sampling sites. The chosen resolution of the grid allowed for most individuals in the same sampling site to be assigned to a unique deme, based on preliminary explorations. We used the cross-validation approach (Marcus et al., 2021) to choose the penalty parameters to be used in the optimization procedure (λ and λ_q_; https://github.com/VivaswatS/feems/blob/main/docsrc/notebooks/cross-validation.ipynb). After preliminary explorations, the migration surface presented here was generated using λ = 0.17 and λ_q_ = 100, which minimized the cross-validation error (although lower values appeared roughly as adequate, Figure 6).

### Analysis of genetic variation across the *bairdi* x *chrysops* hybrid zone

#### Calculation of a genomic hybrid index and genetic composition of the hybrid zone

We estimated a genomic hybrid index and interclass heterozygosity using functions implemented in *triangulaR* (Wiens & Colella, 2025). As noted above, the southern parental population was defined based on our own results and includes individuals from central to south Ecuador that do not cluster genetically with *griseonuchus* (see Results). In the SNP dataset for this analysis, we included all SNPs per locus.

To identify ancestry-informative SNPs, we chose an allele frequency difference of 0.5 between parental populations. Higher thresholds resulted in very few (i.e., less than 100) or no SNPs being retained, and there were no alleles fixed in each parental population. Then, to describe the genetic composition of the hybrid zone, we used the values of genome-wide hybrid indices and observed interclass heterozygosity for all individuals, and plotted these in a triangle plot (Cronemberger et al., 2020; Liu et al., 2022; Lopez et al., 2021).

#### Geographic cline analyses

To describe the geographic extent of the hybrid zone and evaluate whether genetic and plumage variation were concordant, we fitted geographic clines using *HZAR* (Derryberry et al., 2014). We fitted and compared clines for genomic hybrid indices—corresponding to admixture proportions and estimated using *gghybrid*—and plumage hybrid indices estimated previously for these individuals (Céspedes-Arias et al., 2021). For each of these “traits”, we fit a model that estimates cline width (*w*) and center (*c*) but has fixed values for mean and variance of the trait values at the cline extremes (*μL, μR, varL, varR*). Both models were run with a tune value of 1.5, and for each of these we ran 3 chains of 2 million iterations discarding the first 10% as burn-in. All runs were performed randomizing starting parameters. To evaluate whether plumage and genomic clines are coincident and concordant, we compared the 95% confidence interval of the center and width estimates, which were calculated aggregating the three independent chains for each model.

## Results

### Evolutionary relationships among plumage groups as inferred from phylogenies and admixture graphs

We employed two different phylogenetic approaches to describe evolutionary relationships among the seven plumage groups that define the study taxa. Using a maximum-likelihood approach with all loci concatenated (21,304 variant sites), we found that most individuals of the same subspecies clustered together (Figure 1, panel C), yet we did not observe complete reciprocal monophyly in most groups. The *chrysops* group was the exception, as it appeared as a clade nested within the *ornatus* population. Notably, however, support values were generally low, implying that not many sites are phylogenetically informative as expected for a scenario of recent divergence and/or ongoing gene flow. Generally, we observed that individuals from geographically adjacent subspecies are intermixed in the tree.

The *snapper* subspecies-level tree (Figure 2A), inferred using 3,105 biallelic SNPs, showed strong support for a monophyletic “southern” group, encompassing three *M. melanocephalus* subspecies (*malaris, melanocephalus, bolivianus*) characterized by having spectacles and a black crown. The taxon *griseonuchus*, from north of the Marañón River Valley, characterized by having a reddish crown that is phenotypically very similar to *bairdi*, was recovered as sister to a strongly supported clade including the more northerly groups from Colombia and Ecuador. Within this expanded “northern” group, *chrysops* was recovered as sister to *ornatus*, and these, in turn, are sister to *bairdi* (Figure 2A). Note that, unlike the maximum-likelihood tree, we *a priori* assigned individuals to groups for these analyses, which can bias the phylogenetic estimation given that the limits among these populations are not fully defined.

**Figure 2.**
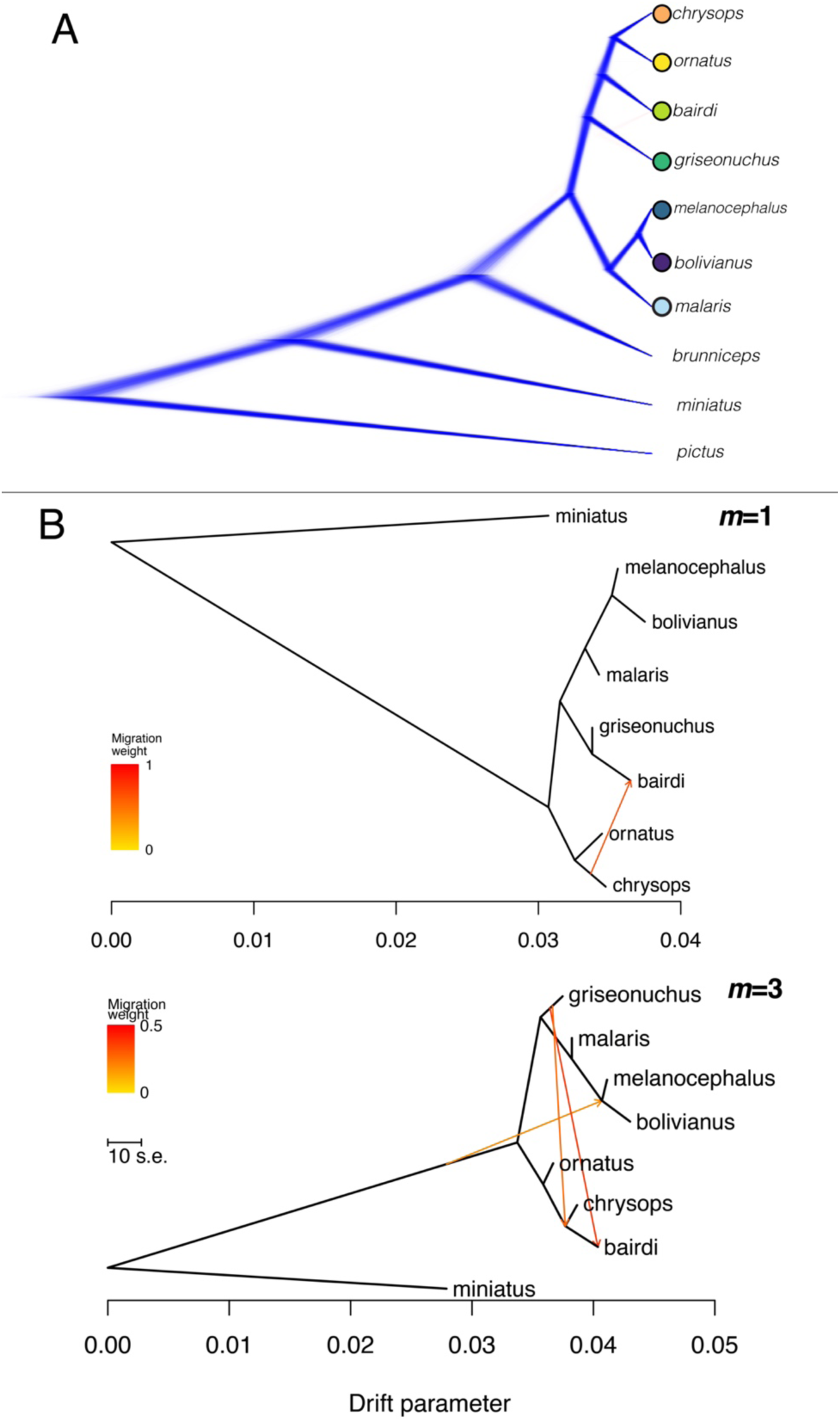
Evolutionary relationships within groups in the *Myioborus ornatus-melanocephalus* complex as inferred by snapper and *OrientAGraph*. **(A)** The main figure shows the phylogenetic relationships between taxa, with the blue topology corresponding to the most frequent in the posterior distribution, followed by red. **(B)** Admixture graphs from *OrientAGraph* with 1(top) and 3 (bottom) migration edges (*m*). The migration edges are shown as colored arrows, and the hue denotes the migration weight.

Overall, we found stronger signals of admixture events between the northern relative to the southern group (Figure 2B). Depending on the *m* value, we recovered slightly different topologies, though we note that these incongruencies were restricted to the relationships between the northern group taxa, namely *griseonuchus, bairdi*, *chrysops*, and *ornatus* taxa (Figure 2B). Here we present the inferences for m = 1 and m = 3 as these were the optimal values based on the linear regressions fitting the likelihood vs *m* curves (Supplementary Figure 1). Both graphs consistently recovered a monophyletic “southern” group (*malaris*, *melanocephalus* and *bolivianus*), which was consistent with the topology of the *snapper* tree. In the m = 1 graph (Figure 2B), *griseonuchus* and *bairdi* appeared as sister, and this clade was, in turn, sister to the southern group. In contrast, in the m=3 graph, *bairdi* was sister to *chrysops*, and thus a migration edge between these taxa was not estimated despite the known migration event parameter. Two edges with high migration weight were inferred, between *griseonuchus* and *bairdi,* and *griseonuchus* and the *bairdi-chrysops* node. An additional edge, with low migration weight, was inferred between the outgroup and the *bolivianus-melanocephalus* group (Figure 2). We note, however, that the outgroup has a very low sample size (n = 3) and thus do not find this inference to be very reliable.

### Genetic structure within the *ornatus-melanocephalus* complex

The PCA, based on 4,800 biallelic SNPs, showed that individuals generally cluster in relation to plumage groups and geographic origin (Figure 3), but some phenotypically distinct groups were very close in the PC1-2 space (particularly *chrysops, ornatus* and *bairdi*). As in the phylogenetic analyses, one can distinguish clearly the “southern” group, including all individuals from populations south of the Marañón River Valley (*bolivianus, melanocephalus, malaris,* Figure 3), which occupied a wide range of PC1 values (Figure 3). There was also an identifiable northern cluster (*chrysops, bairdi*, and hybrid zone individuals) that was, however, not fully distinct from a third cluster comprised of *griseonuchus* individuals. Within this northern cluster, the plumage groups *ornatus, chrysops, bairdi,* and individuals from the hybrid zone (“northern”, hereafter) appeared very close in PC1-2 space. As expected, individuals from the hybrid zone (in gray, Figure 3) fell intermediate between their parental populations. The *ornatus* group appeared as a more defined cluster when considering PC3 (Supplementary Figure 5). Notably, some individuals putatively classified as *bairdi*, collected in the Ecuadorian provinces of Loja and outlining ridges in the Amazon, appeared to cluster with *griseonuchus* (Figure 2). We observed a similar pattern in the hybrid zone plus *griseonuchus* PCA (Supplementary Figure 6). We obtained similar results when using a SNP set of the whole complex obtained with a stricter filtering for the PCA (Supplementary Figure 7).

**Figure 3.**
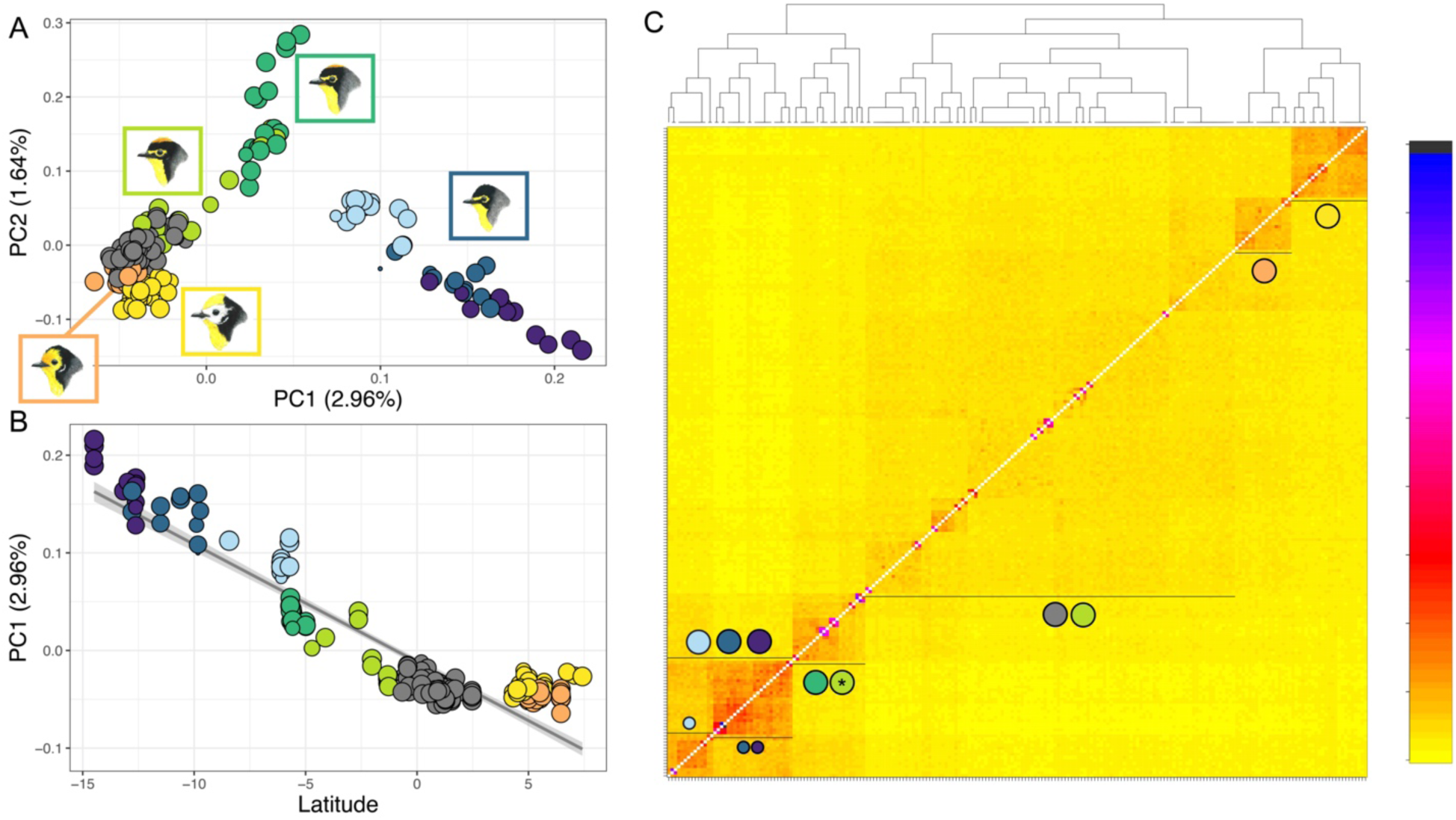
Genetic structure within the *ornatus-melanocephalus* complex. (**A)** The distribution of individuals in the PC1/PC2 space showing clustering based on plumage groups, which are denoted by the color of the dots. **(B)** Strong correlation between PC1 and latitude. In both cases (A and B) the size of the dot is inversely proportional to the proportion of missing data. **(C)** fineRADStructure coancestry matrix, and associated distance tree (above) showing clusters of individuals that overall correspond to plumage groups, which are denoted by the colored dots that correspond to groups as in Figure 1. The asterisk highlights a small number of putatively *bairdi* individuals, all from the south end of Ecuador, that genetically cluster with *griseonuchus* (from northern Peru). The color scale denotes values of coancestry, with blue and black tones corresponding to very high levels of coancestry.

Two main clusters are supported in the coancestry matrix resulting from *fineRADstructure*: populations occurring from south Ecuador to Bolivia (*griseonuchus* plus*“*southern” group), and those occurring from Ecuador to Colombia (“northern” group) (Figure 3C), which contrasted with the topology inferred by *snapper*. Within the first group, *griseonuchus* and the southern group appear as differentiated clusters. The few exceptions to this pattern are a few putatively *bairdi* individuals that clustered with *griseonuchus*, but were collected in the southern edge of the distribution of this plumage group (where it is replaced by *griseonuchus,* Figures 1 and 3) in the Cordillera del Cutucú and Loja Province of Ecuador. We detected two main groups within the “northern” group (Ecuador and Colombia): one corresponded to *ornatus* (occurring in the Eastern Andes of Colombia) and the other to the hybridizing taxa in the complex and individuals from the hybrid zone.

Admixture analyses including all individuals in the *ornatus-melanocephalus* complex revealed genetic breaks along the distribution of these warblers that were overall consistent with fineRADStructure (Figure 4, A). For example, when assuming K = 2 and K = 3 the “southern” group appeared as an identifiable cluster. For K = 2, which is the optimal K value based on cross-validation error values (Supplementary Figure 8, top panel), the group *griseonuchus* also clustered with the southern group (Figure 4, A1). For K = 3, a “southern” plus *griseonuchus* group can be also identified, and there was an additional genetic break roughly located in the described *bairdi-chrysops* hybrid zone in southern Colombia (Figure 4, B1).

**Figure 4.**
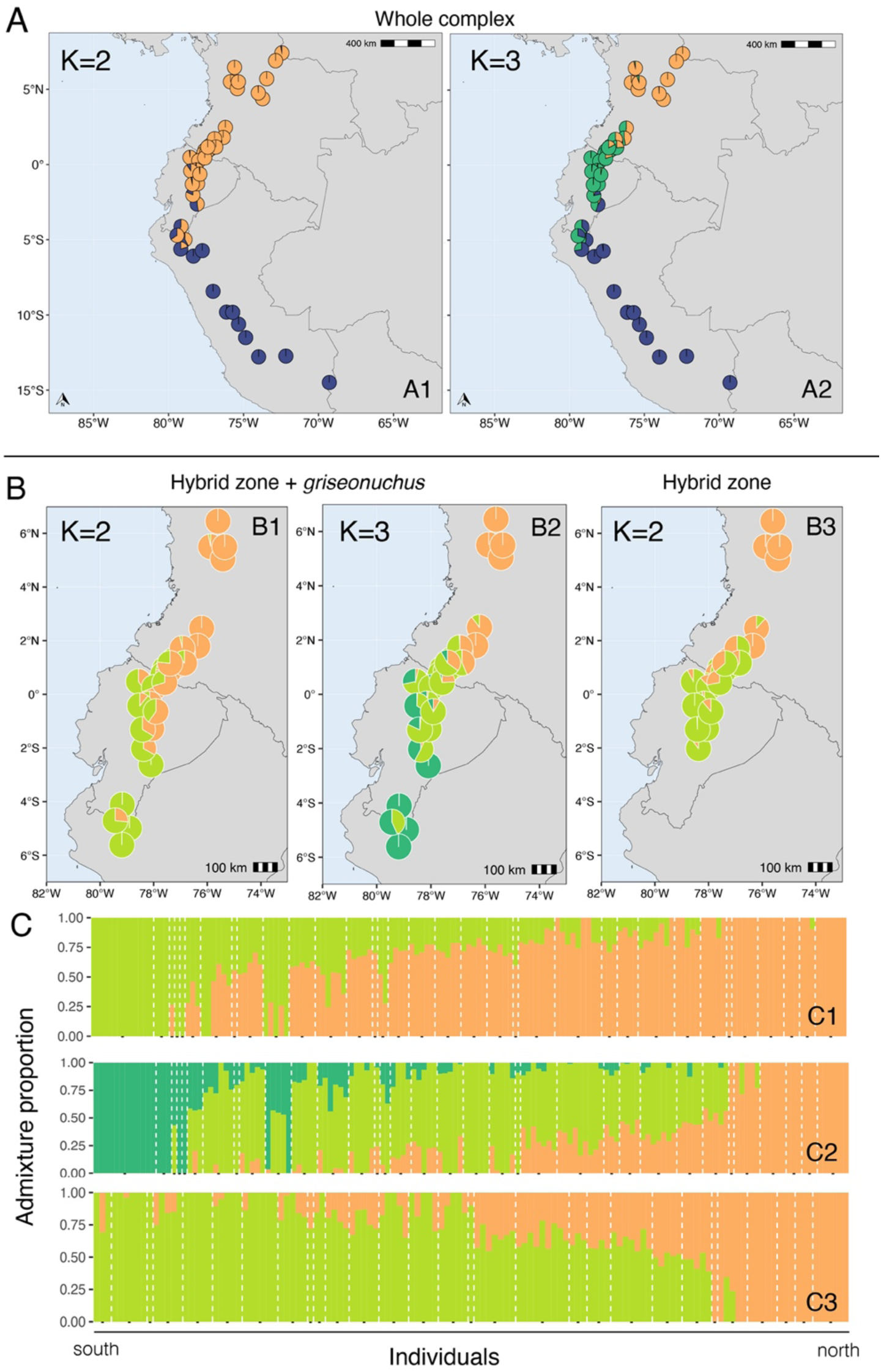
Genetic clusters (**A**) across the entire distribution of the *ornatus-melanocephalus* complex and (**B**) the hybrid zone and adjacent populations as shown by the mean admixture proportions per locality. The dataset and *K* value for each plot is specified at the top of each map. **(C) Admixture prop**barplots showing the admixture proportions per individual, which are organized into localities from south to north (localities are divided by white dotted lines).

The results of an admixture analyses focused on the hybrid zone and adjacent populations, when assuming K = 2, showed a genetic break that clearly coincides with the *chrysops-bairdi* hybrid zone (Figure 4, B and C), as previously identified based on plumage variation (see map in Figure 1). However, when also considering *griseonuchus*, the transition between these genetic groups is less sharp and displaced to the south (Figure 4, panel B). In both cases there was some east-west differentiation, with localities to the west of the Ecuadorian Andes showing a higher mean admixture proportion corresponding to the southern cluster (Figure 4, B1 and B3). When using K = 3 for the analyses including *griseonuchus* (a *K* that corresponds to the number of identifiable plumage groups), two genetic clusters were identified in Ecuador, and one in Colombia, and one of these genetic breaks did again roughly correspond to the *bairdi-chrysops* hybrid zone (Figure 4, B2). In both cases the *K* value with the lowest cross-validation is *K* = 1 (Supplementary Figure 7).

Values of pairwise F_ST_ between populations showed an overall correspondence with geographic distance: subspecies that are far apart in space show higher genetic differentiation (e.g., *ornatus* vs *bolivianus,* Figure 5). We observed higher pairwise differentiation between taxa belonging to the “southern” versus “northern” groups, which is consistent with all other analyses describing genetic structure. In contrast, there was very low differentiation among the three subspecies occurring south of the Marañón River Valley (“southern” group), which all have similar head coloration. However, there was also relatively low levels of differentiation within the “northern” plus *griseonuchus* group, which encompasses striking plumage diversity, particularly between the two hybridizing taxa, *bairdi* and *chrysops, and bairdi* and *griseonuchus* (Figure 5). There was a positive correlation (Figure 5, Mantel statistic r = 0.75, p-value = 0.001) between pairwise F_ST_ calculated based on our ddRAD SNPs and the ND2 mitochondrial gene (Céspedes-Arias et al., 2021). Lastly, as expected if patterns of genetic variation reflect IBD, we found a positive correlation between pairwise F_ST_ between localities and geographic distance (Figure 5, Mantel statistic r = 0.65, p-value = 0.001). Using the same classification scheme for individuals as that to calculate F_ST_, we estimated genetic diversity statistics. We found that nucleotide diversity is very similar among groups, ranging from 0.076 to 0.091 (Supplementary Table 3).

**Figure 5.**
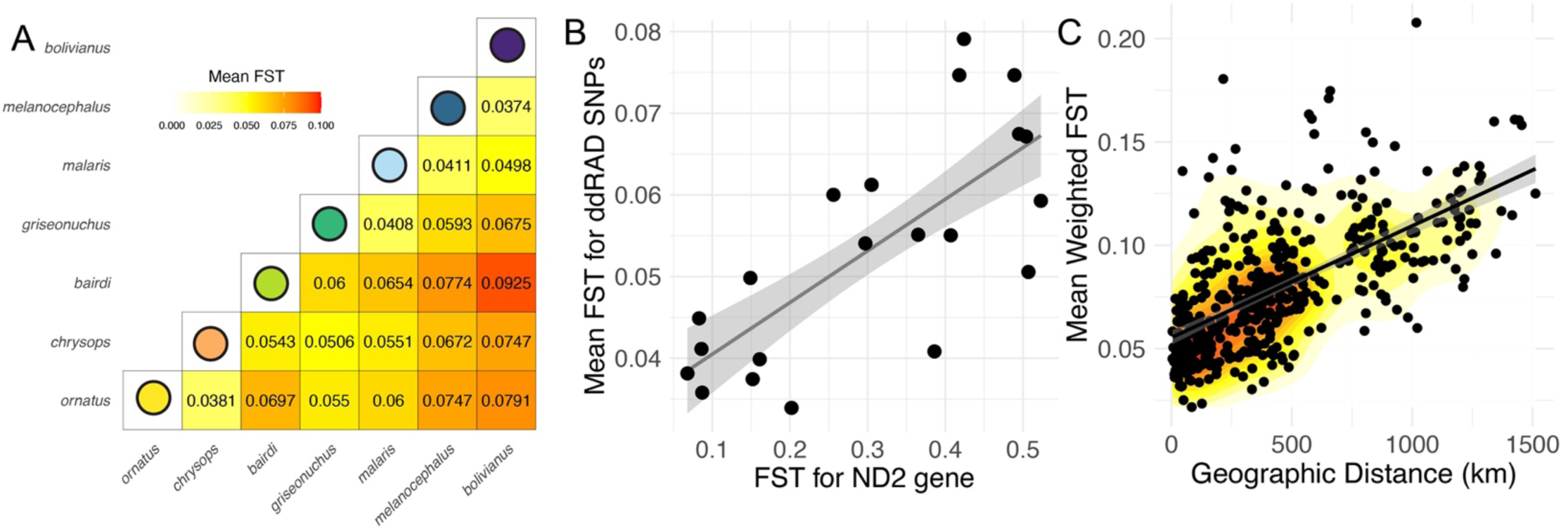
Genetic differentiation among Andean *Myioborus* taxa. **(A)** Mean pairwise FST values between subspecies (i.e., plumage and geographic groups). **(B)** Correlation between pairwise FST calculated with ddRAD SNPs (this study) and the ND2 mitochondrial gene from the same set of samples (Cespedes-Arias *et al* 2021). **(C)** Positive correlation between geographic distance and FST between localities supports a role of isolation-by-distance shaping genetic variation in the *ornatus-melanocephalus* complex.

The migration surfaces inferred by FEEMS suggest that there are areas within the distribution of these warblers in which genetic differentiation cannot be fully explained by IBD (i.e., areas of inferred low gene flow; Figure 6). These areas of low migration coincided mostly with two geographic breaks that dissect the distribution of the complex (Figure 1, Figure 6), specifically the Magdalena River Valley and the Marañón River Valley, and with the hybrid zone. In contrast, most of the range south of the North Peruvian low showed mid to high values of inferred effective migration, which can indicate that genetic structure along this area is well explained by IBD alone. We acknowledge that low sampling density in certain regions, particularly in central and southern Peru, may have influenced the inferred migration surface (Figure 6). However, cross-validation results indicate that the selected regularization parameter provides a good fit while accounting for unobserved nodes (Supplementary Figure 9). With that caveat in mind, the FEEMS results are broadly consistent with the patterns of genetic structure revealed by our other analyses.

**Figure 6.**
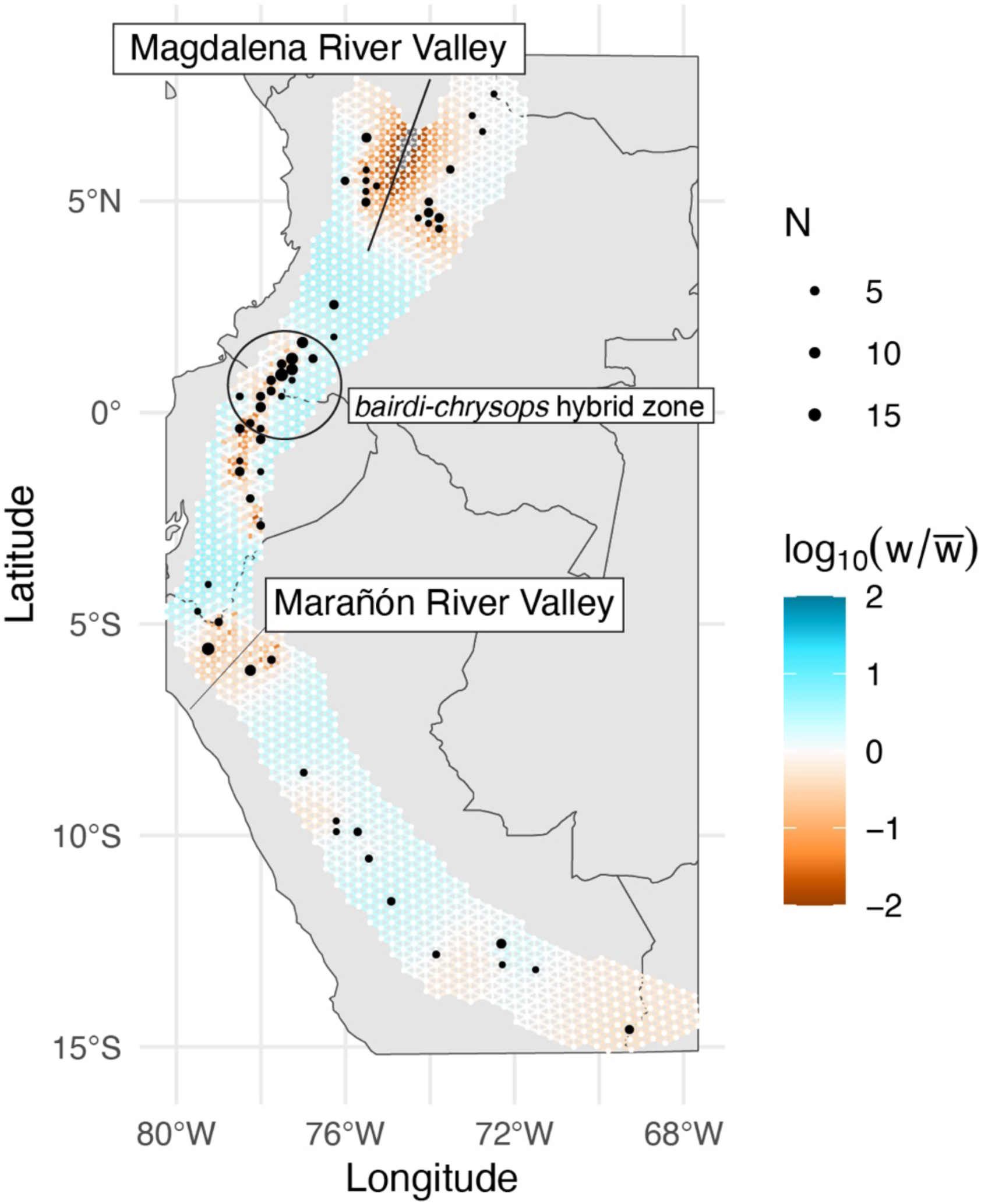
Estimated effective migration graph across the entire distribution of the *Myioborus ornatus-melanocephalus* complex, highlighting areas of restricted gene flow. The edges of the grid are colored by estimated effective migration, from burnt orange (low migration) to blue (high migration). In the grid, the black dots correspond to the sampled nodes and the size of the dot correspond to the sample size (*N*). We highlight three areas of low migration, two geographic barriers and the hybrid zone where genetically distinct taxa have come into contact.

### Hybrid zone

To describe the geographic extent of hybridization and the genetic composition of the *bairdi* x *chrysops* hybrid zone, we identified ancestry-informative markers to estimate genomic hybrid indexes and interclass heterozygosity for individuals across the hybrid zone and parental *bairdi* and *chrysops* populations. Using a relatively low threshold for allele frequency difference (0.5), we identified only 29 sites that were ancestry informative from the total of 14,301, and none of these were fixed between the parental populations. This result complicates the interpretation of the triangle plots (see below) and highlights the low level of genetic differentiation between these two hybridizing taxa. Because of the scarcity of ancestry informative sites in our ddRAD data set, which can affect the accuracy of genomic hybrid index estimations, we opted to conduct cline analyses with admixture proportions (with K=2) instead.

We found that the head coloration and genomic hybrid index, defined as the admixture proportions in Figure 4 (panel C3) had different centers but similar widths (Figure 7, panel A). The admixture proportion cline had a width of 281.6 km (95% confidence interval [CI]: 245.0‒ 319.8km), compared to 240.6 km for head coloration (95% confidence interval [CI]: 277.5‒311.1 km). Given the substantial overlap in confidence intervals, we do not interpret this 41 km difference in width as significant. However, we found a 37 km difference in the estimates of the center between the plumage and admixture proportion clines, for which the 95% confidence intervals did not overlap. For admixture proportions the cline was centered around 348.8 km (95% confidence interval [CI]: 335.4‒362.7 km) in our sampling transect, while the plumage cline was centered around 311.1 km (95% confidence interval [CI]: 300.7‒322.6 km). Therefore, based on our *HZAR* analyses, the plumage cline appeared to be displaced towards the south (i.e., towards Tungurahua, Ecuador, the origin of the sampling transect), implying that the *chrysops*-like plumage traits may be introgressing into *bairdi*. However, there was an overall strong correlation between the plumage hybrid indices and admixture proportion across the hybrid zone and allopatric populations (Figure 7, panel B; linear model; *p-value* < 0.001, *m* = 0.9, *R-squared* = 0.8). We identified the outliers of this linear regression as those individuals with residuals >2 SD units from the mean, finding that most were below the regression line (Figure 7). Specifically, we identified six individuals that had a more *chrysops*-like plumage from the hybrid zone area (i.e., lower) relative to what the model predicts based on the admixture proportion, which was consistent with the cline analysis results that shows that *chrysops*-like plumage is introgressing into *bairdi*.

**Figure 7.**
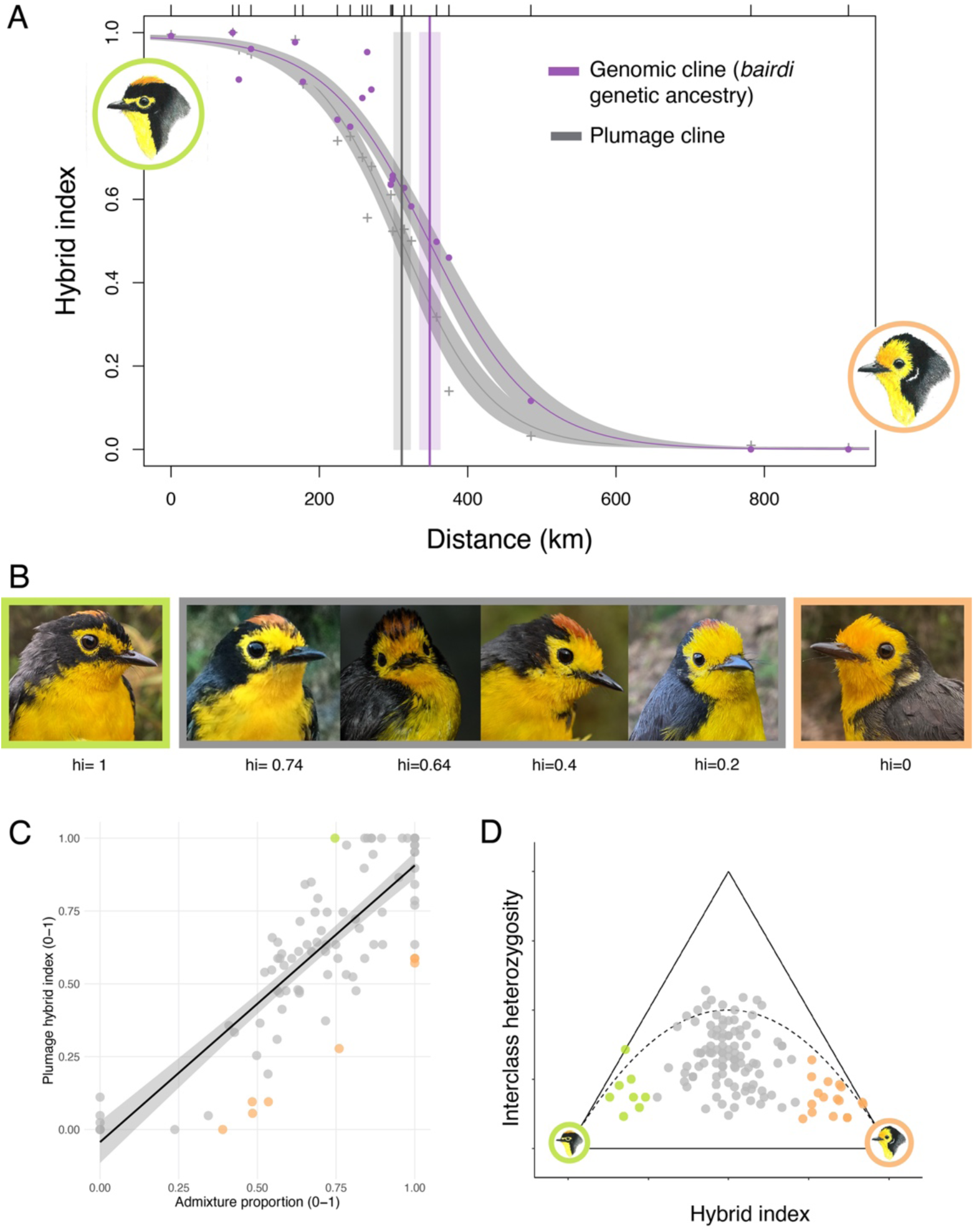
Genetic and plumage variation across the *bairdi x chrysops* hybrid zone. **(A)** The plumage and genomic clines have different estimated centers. **(B)** Photographs of specimens collected for which we calculated a plumage hybrid index, showing a sample of the phenotypic diversity across the hybrid zone, and “parental” phenotypes at both extremes. Photos: Andrés Cuervo, Paulo Pulgarín, David Ocampo, and Laura Céspedes. **(C)** We observed an overall a strong correlation between the genomic hybrid index (*bairdi* admixture proportion) and the plumage hybrid index. Most individuals that deviate from this relationship (outliers marked in color) are individuals with a more chrysops-like plumage than expected given the admixture proportion. (**F)** Triangle plot showing the genetic composition of the hybrid zone. The x axis corresponds to a genomic hybrid index as calculated using 29 ancestry informative markers. Parentals are shown in lime green and orange.

To describe the genetic composition of the *bairdi-chrysops* hybrid zone, we built triangle plots (interclass heterozygosity vs hybrid index) based on the 29 ancestry-informative markers that we identified (Figure 7, panel C). The individuals from the hybrid zone are distributed towards the center of the triangle, many below the line that defines what value combinations are possible under Hardy-Weinberg Equilibrium which is likely a product of the low allele frequency difference threshold. We note that individuals from populations away from the hybrid zone (Figure 7, in green and orange) are not located in the bottom corners of the triangle plot, which is expected given that the allele frequency difference threshold is lower than one. If interpreted considering the caveats described above, the distribution of individuals in the triangle is suggestive of a hybrid zone mostly composed of advanced generation hybrids, without many F1s.

## Discussion

We used reduced-representation genomic data to study genetic variation at two different spatial scales: the full distribution of a geographically variable warbler complex (*M. ornatus-melanocephalus*) and a known hybrid zone between two of these plumage forms. At both scales, our set of thousands of SNPs allowed us to explore genetic structure with a resolution that mtDNA loci could not provide in previous studies (Céspedes-Arias et al., 2021; Pérez-Emán, 2005). We found evidence of isolation-by-distance driving genetic structure across the latitudinally extensive range of these warblers, and genetic divergence increased where ranges are separated by known geographic barriers. These clear genetic breaks coincided with observed differences in head plumage coloration among populations. Notably, one transition between groups includes a geographically restricted hybrid zone between *bairdi* and *chrysops*, likely comprised primarily of advanced generation hybrids, where differences in genetic ancestry and head coloration are correlated.

Below we discuss how our findings relate to the history of divergence and gene flow among these populations, with a focus on how genetic variation imperfectly covaries with plumage traits, and how it is structured across geography. We then discuss how genetic variation is structured across the hybrid zone, how it relates to patterns of variation in head coloration in this area, and whether there is evidence of selection against hybrids maintaining this seemingly extensive hybrid zone.

### Geographic variation in plumage color and genetic structure

Our qualitative assessments based on historical plumage descriptions and examinations of study skins suggest that for these *Myioborus*, patterns of geographically clustered phenotypic variation coincide with phylogeographic structure uncovered in the present study. First, we found that individuals belonging to the same subspecies usually form genetic clusters, and that all major genetic breaks within the species complex coincide with a transition between clearly differentiated plumage groups, although we note that we lack data quantifying the degree of plumage differentiation between subspecies (e.g. Winger and Bates 2015). In addition, subspecies with subtle plumage differentiation (Cuervo and Céspedes Arias 2023), specifically those occurring south of the Marañón River Valley (Figure 1), all characterized by black crown and spectacles, show little genetic differentiation from one another.

Previous examinations of genetic variation in mitochondrial loci (Céspedes-Arias et al., 2021; Pérez-Emán, 2005) found extensive haplotype sharing and little genetic differentiation among the subspecies *ornatus* and the hybridizing *bairdi* and *chrysops*, which are all characterized by striking plumage differences (see Figure 1). In contrast, in our analyses of ddRAD data, which encompass thousands of SNPs across the genome, these subspecies are each identifiable genetic clusters, albeit with shallow genetic differentiation. Taken together, we interpret these results as evidence of rapid and recent diversification of head plumage patches in the *ornatus-melanocephalus* complex, as seen in other Neotropical taxa (Campagna et al., 2012, 2017). Our results also highlight the usefulness of reduced representation genomic data to study the evolutionary history of young radiations (Aguillon et al., 2018; Toews, Campagna, et al., 2016), in which few loci might be insufficient to detect genetic differentiation and describe evolutionary relationships among phenotypically distinct populations (Friis et al., 2016; Stryjewski & Sorenson, 2017). We note that lack of differentiation in mtDNA between phenotypically distinct groups could also be the result of mitochondrial introgression (Irwin et al., 2009; Toews & Brelsford, 2012), although the strong positive correlation of F_ST_ for ND2 and our ddRAD loci does not support a scenario of extensive mitochondrial introgression in the group.

Genetic structure in the region of the Andes spanning from central Ecuador to northern Peru (north of the Marañón River Valley) is complex and not clearly associated with a single topographic feature. This area also broadly corresponds to the transition between *griseonuchus* and *bairdi* plumage (see Figure 1, Supplementary Figure 1), two subspecies characterized by a rufous crown and spectacles. Despite their overall similarity, these two groups show differentiation in at least two head plumage traits: the presence of a black band behind the rufous crown and the extent of black in the cheeks and lores (Zimmer 1949; Curson et al. 1994; World and International 2019; Cuervo and Céspedes Arias 2023). Our qualitative assessments of plumage variation (Supplementary Figure 1), however, suggest that the variation in these traits might not be completely discrete. Moreover, we found that individuals from Loja and the Cordillera del Cutucú (Ecuador), show substantial variation in both putatively diagnostic plumage traits. Therefore, these individuals, which are genetically clustered with *griseonuchus,* do not uniformly resemble the individuals from Cajamarca, Peru (Supplementary Figure 1). Because of limited sampling in the area, we are unable to draw stronger conclusions, but our data suggest that there is some discordance between plumage and genetic variation, and evidence of introgression based on admixture graphs. Despite some differentiation in at least these two plumage traits, our genetic results are concordant with substantial historical or contemporary gene flow among populations in this area. This suggests that the geographic range of *bairdi* is potentially outlined by two hybrid zones, one to the north with *chrysops* (Céspedes-Arias et al., 2021), and to the south with *griseonuchus.* Evaluating this hypothesis of hybrid zones at the range edges, as observed in barn swallows (Scordato et al., 2017), will require a systematic collection of specimens across the *bairdi-griseonuchus* boundary.

### Genetic structure across space: the influence of geographic distance and topographic features along the Andes

Several of our analyses support a strong correlation between genetic variation and geographic location of samples and support an important role of isolation-by-distance driving genetic structure in *Myioborus* along the Andes. Patterns of IBD are close to ubiquitous in nature (Aguillon et al., 2017; Seeholzer & Brumfield, 2018; Vos & Velicer, 2008), but see Meirmans, 2012), and are often taken as the null model for genetic differentiation if organisms have limited dispersal given the scale of their range, as is the case for these small warblers with a latitudinally-expansive distribution range. However, IBD does not account fully for the observed patterns of genetic variation. Specifically, we found evidence of discrete genetic—and phenotypic—clusters, and of reduced migration (relative to IBD) across some areas along the Andes, which in turn correspond to lowland valleys such as the Marañón and the Magdalena River Valleys. These deep, lowland gaps interrupt the continuity of humid high-elevation forest and scrub (Graham et al., 2010) that these *Myioborus* inhabit in the present, and also coincide with genetic and phenotypic breaks within other taxa that inhabit similar elevation bands (Gutiérrez-Pinto et al., 2017; Martínez-Gómez et al., 2023; Winger & Bates, 2015). These two topographic features were identified in a comparative study as being associated with allopatric events across multiple Andean bird genera (Hazzi et al., 2018). These genetic breaks, clearly tied to topographic features, also correspond with striking discontinuities in head coloration traits (see Figure 1).

Other areas associated with clear genetic breaks observed in the inferred migration surface were not clearly associated with topographic features that interrupt the present-day continuity of humid, high<u>-</u>elevation forests. Specifically, low migration is inferred in a large area ranging from the border between Ecuador and Colombia down to southern Ecuador. We suggest that this pattern emerges results partially because this broad area includes the *chrysops-bairdi* hybrid zone, in which genetic differentiation relative to distance is expected to be higher than what is expected under pure IBD, as two somewhat differentiated subspecies come into contact. The extension of this area of low inferred migration to south Ecuador is consistent with our fineRADStructure, PCA and admixture results that suggest that there is substantial genetic differentiation between individuals in close proximity in this area. Specifically, we found that birds from isolated mountain spurs such as the Cordillera del Cutucú (southeastern Ecuador) and Cordillera del Condor (Peru-Ecuador border) clustered genetically with birds from the main Andes in Cajamarca (northwestern Peru), and Loja (southern Ecuador), instead of the more geographically proximate individuals from the eastern Ecuadorian Andes in Morona-Santiago (see Figure 2). The Cordillera del Cutucú and Cordillera del Cóndor are outlying mountain ranges in the Amazonian foothills, separated from the main Andes by the valleys of the rivers Upano and Zamora, respectively (Pozo-Zamora et al., 2022; Robbins et al., 1987). Our results suggest that these valleys, and surrounding lowland areas, have likely acted as dispersal barriers for these higher-elevation warblers. On the other hand, the patterns of genetic variation described here indicate that there has likely been substantial gene flow connecting the populations of warblers inhabiting isolated peaks in these mountain ranges (separated by the Santiago River Valley; Robbins et al. 1987; Pozo-Zamora et al. 2022), and the eastern slope of the Andes in northern Peru.

### Hybrid zone dynamics

Based on our reduced-representation genomic data set, the hybridizing *chrysops* and *bairdi* are characterized by very shallow genetic divergence, which contrasts with their striking plumage differentiation. Moreover, across thousands of SNP sites, we only identified 29 ancestry informative sites with substantial allele frequency differences between parental populations. This scarcity of ancestry informative markers limits our ability to draw definitive conclusions about the genetic composition of this hybrid zone but highlights the low level of genetic differentiation between *chrysops* and *bairdi*.

This hybrid zone likely formed by secondary contact of two formerly isolated plumage forms. The disconnection and subsequent reconnection of habitat that could have shaped this process may have followed elevational shifts in the cloud forest belt during the Pleistocene (Flantua et al., 2014; Ramírez-Barahona & Eguiarte, 2013), or potentially been driven by volcanic activity that modified topographic connectivity (Sanín et al., 2024). An investigation of the evolutionary origins of this hybrid zones will require both a better understanding of the patterns of connectivity of cloud forest in the area where the *chrysops-bairdi* hybrid zone is centered, as well as an estimation of the time since secondary contact, which could be obtained based on inferences of ancestry tract length in whole genome data (Pool & Nielsen, 2009).

Our geographic cline analyses using plumage and genomic data suggest that the hybrid zone extends for over 200 km, similar to the findings from its original description based on face color traits (Céspedes-Arias et al., 2021). We do not find statistically significant differences in the width of the admixture proportion (i.e., genomic) and plumage geographic clines. Taken together, plumage and genomic patterns of variation thus suggest that hybridization between *chrysops* and *bairdi* has been extensive, albeit still restricted to a fraction of the entire distribution of these taxa (approximately 30% of our sampling transect) which is continuous at least from central Ecuador to the north of the Central Andes of Colombia. Therefore, the geographic extent of hybridization is unlikely to be constrained by physical or ecological barriers that limit dispersal beyond the hybrid zone (e.g., DeRaad et al., 2023), raising the question of what factors are restricting the hybrid zone to this area.

The lack of information on natal dispersal distances as well as a the wide range of plausible times since secondary contact constrains our ability to generate a direct comparison of the observed cline widths with that expected under neutral diffusion (Barton and Gale 1993). With the available data we are thereby unable to evaluate whether the *chrysops* x *bairdi* hybrid zone corresponds to a stable tension zone, in which the width of clines is constrained by selection against hybrids. A second possibility is that the observed sharp genomic and plumage clines are not maintained by selection, but instead that this is a young and potentially expanding hybrid zone, and that the differentiation between *chrysops* and *bairdi* might become eroded over time (Kearns et al., 2018).

Using a range of plausible dispersal distances inferred from another *Myioborus* species (Céspedes-Arias et al., 2021; Mumme, 2015), we can approximate expectations under neutrality (Barton & Gale, 1993). For a cline width of 281 km, the neutral diffusion model predicts that ∼14,337 generations would be required assuming a dispersal distance of 935 m, or ∼53,278 generations with a dispersal distance of 485 m. In principle, building upon our systematic collection of specimens across the *chrysops* x *bairdi* hybrid zone, the alternative of an expanding hybrid zone could be revisited by comparing plumage and genomic clines based on specimens collected across different time points. However, given that the expansion of this hybrid zone under neutral diffusion with plausible dispersal distances is expected to be slow (i.e. 0.5 km over 50 generations), time between sampling periods would need to be very long to detect changes.

We found that cline centers are not fully coincident, as that of the plumage cline is displaced to the south of the genomic cline center, implying that the *chrysops*-like plumage is introgressing into *bairdi.* Assuming that most of our ddRAD loci can be considered neutral markers, our results indicate that head coloration might be under selection across this hybrid zone (Gay et al., 2008). Specifically, the observed pattern would be consistent with a scenario in which the *chrysops*-like plumage is favored across the hybrid zone, which would lead to the asymmetric introgression of this phenotype towards the south. Asymmetric introgression of plumage traits has been observed in other avian hybrid zones (Baldassarre et al., 2014; Lim et al., 2024; Semenov et al., 2021), and in two of these cases, behavioral evidence supported the idea that these traits provide fitness advantages in the context of mate choice (Baldassarre & Webster, 2013; Long et al., 2024; Stein & Uy, 2006).

Even though the difference in plumage and genomic cline center is statistically significant, we found it to be relatively small (38 km; approximately 4% of the entire sampling transect). This contrasts with much more marked differences in center estimations found in other hybridizing birds like *Motacilla* wagtails (Semenov et al., 2021) and *Malurus* fairy-wrens (Baldassarre et al., 2014). We acknowledge that this minor difference could be an artifact of the way in which both of our hybrid indexes were calculated (e.g. with the scoring system giving equal weight to pre-defined patches; Céspedes-Arias et al., 2021), and potentially low sampling sizes in some of the localities. However, this slight difference could be the result of selection on the head plumage patches. Even though the role of these color patches has not been studied in these warblers yet, it is plausible that they may be relevant for intraspecific communication and mate choice as seen in head plumage patterns in many other bird species (e.g., Marchetti 1992; Andersson et al. 1998; Scott and Deag 1998; Rémy et al. 2010). Future behavioral studies could evaluate these hypotheses of selection favoring *chrysops*-like head color patterns across this hybrid zone.

Besides describing the geographic extent of hybridization, we used our reduced-representation genomic data set to gain insights into the genetic composition of the *chyrsops x bairdi* hybrid zone. However, as noted above, the scarcity of ancestry informative sites in our dataset complicates the interpretation of these results, as it compromises the accuracy of genomic hybrid indexes and interclass heterozygosity estimations (Gompert et al., 2024; Wiens & Colella, 2025). Setting a low allele frequency threshold, which was necessary in our case, can also result in a tendency of individuals to cluster near the center of triangle plots (Wiens & Colella, 2025). Bearing these caveats in mind, we interpret our triangle plot results as suggestive of a hybrid zone composed mostly of advanced generation hybrids, with no F1s sampled, given the lack of individuals with intermediate hybrid indexes and very high interclass heterozygosity (Capblancq et al., 2020; DeRaad et al., 2023; Natola et al., 2021). This is consistent with the existence of a high diversity of plumage phenotypes in the hybrid zone, as well as the lack of contemporary geographic contact between parental *chrysops* and *bairdi* populations (i.e., we do not observe “pure” individuals of both species in the same sites) (Céspedes-Arias et al., 2021). Given the shallow genetic divergence between *chrysops* and *bairdi*, a more detailed examination of the genetic composition of this hybrid zone will necessitate whole-genome sequencing, capturing more ancestry informative sites. Specifically, as in other examples of hybridizing taxa with very little genomic divergence, whole-genome sequencing may be needed to identify these regions and sites that are highly differentiated and study hybridization dynamics in detail (Aguillon et al., 2021; Poelstra et al., 2014; Toews, Taylor, et al., 2016).

## Author contributions

L.N.C.A, A.M.C., C.D.C. and L.C. conceived the study; L.N.C.A. and A.M.C. conducted fieldwork with substantial logistical support from C.D.C. and E.B.; L.N.C.A. conducted lab work with support from L.C. and I.L.; L.N.C.A. conducted the bioinformatic processing of data and data analyses with substantial input from L.C. L.N.C.A. wrote the original draft, with major contributions from L.C. All authors contributed to reviewing and editing the manuscript.

## Acknowledgements

We are grateful to Bronwyn Butcher for supporting our work at the Fuller Evolutionary Biology Lab (Cornell Lab of Ornithology). We thank the following people for contributing to fieldwork across the *bairdi-chrysops* hybrid zone which resulted in the collection of most specimens included here: P. Montoya, D. Céspedes-Arias, P. Pulgarín, F. González, E. Obando, M.A. Meneses, S. Herrera, P. Cardozo, and D. Ocampo. Our fieldwork was made possible by many landowners, local guides, nature reserve officials, and other members of local communities who allowed us access to their land, contributed with invaluable logistical support and even welcomed us warmly into their homes. We are grateful to the Universidad de los Andes, particularly Mireya Osorio, and to the Universidad Tecnológica Indoamérica for their crucial support in obtaining export permits. Collecting and export from Ecuador (Contrato de Acceso a Recursos Genéticos MAE-DNB-CM-2015-0017 and N° 40-2018-EXP-CM-FAU-DNB/MA to the Universidad Tecnológica de Indoamerica); Colombia (Number 01374 to the Universidad de los Andes). See Cespedes *et al* 2021 for information on collecting permits.

We are also grateful to museum collections that provided tissue loans to conduct this study: Museum of Southwestern Biology, Kansas University Natural History Museum, Louisiana State University Museum of Natural Science, Pontificia Universidad Católica del Ecuador, Instituto de Investigación de Recursos Biológicos Alexander von Humboldt, Universidad de los Andes, and the Cornell University Museum of Vertebrates. We thank Nathan Rice, Jason Weckstein (ANSP), Sebastian C. Perez, Nicholas Mason (LSUMZ) and Santiago Burneo (QCAZ) for providing photographs of museum specimens. Finally, we thank Shannon Hackett, John Bates, Ben Marks (FMNH), Socorro Sierra and Gustavo Bravo (IavH) for facilitating the examination of study skins.

For computing cluster access and technical support, we thank the High-Performance Computing center at Cornell University (BioHPC) and The University of Chicago Center for Research Informatics. We thank Josie Paris for general advice on Stacks use, and Stepfanie Aguillon for suggestions on clustering and phylogenetic analyses. For helpful discussions L.N.C.A thanks the Lovette lab group, as well as Abhimanyu Lele, Louise Bodt, John Bates, Trevor D. Price, Marcus Kronforst, John Novembre and Pavitra Muralidhar. Lastly, we are very grateful to Andrés Montes Rojas, who kindly provided the original illustrations used in the figures.

## References

Aguillon, S. M., Campagna, L., Harrison, R. G., & Lovette, I. J. (2018). A flicker of hope: Genomic data distinguish Northern Flicker taxa despite low levels of divergence. Auk, 135(3), 748–766. 10.1642/AUK-18-7.1

Aguillon, S. M., Fitzpatrick, J. W., Bowman, R., Schoech, S. J., Clark, A. G., Coop, G., & Chen, N. (2017). Deconstructing isolation-by-distance: The genomic consequences of limited dispersal. PLoS Genetics, 13(8), 1–27. 10.1371/journal.pgen.1006911

Aguillon, S. M., Walsh, J., & Lovette, I. J. (2021). Extensive hybridization reveals multiple coloration genes underlying a complex plumage phenotype. Proceedings of the Royal Society B: Biological Sciences, 288(1943). 10.1098/rspb.2020.1805

Alexander, D. H., Novembre, J., & Lange, K. (2009). Fast model-based estimation of ancestry in unrelated individuals. Genome Research, 19(9), 1655–1664. 10.1101/gr.094052.109

Andersson, S., Ornborg, J., & Andersson, M. (1998). Ultraviolet sexual dimorphism and assortative mating in blue tits. Proceedings of the Royal Society of London B: Biological Sciences, 265(November 1997), 445–450.

Andrews, S. (2010). FastQC: A quality control tool for high throughput sequence data.

Baiz, M. D., Wood, A. W., Brelsford, A., Lovette, I. J., & Toews, D. P. L. (2021). Pigmentation Genes Show Evidence of Repeated Divergence and Multiple Bouts of Introgression in *Setophaga* Warblers. Current Biology, 31(3), 643–649.e3. 10.1016/j.cub.2020.10.094

Baldassarre, D. T., & Webster, M. S. (2013). Experimental evidence that extra-pair mating drives asymmetrical introgression of a sexual trait. Proceedings of the Royal Society B: Biological Sciences, 280(October), 1–7. 10.1098/rspb.2013.2175

Baldassarre, D. T., White, T. A., Karubian, J., & Webster, M. S. (2014). Genomic and morphological analysis of a semipermeable avian hybrid zone suggests asymmetrical introgression of a sexual signal. Evolution, 68(9), 2644–2657. 10.1111/evo.12457

Barker, K., Burns, K. J., Klicka, J., Lanyon, S. M., & Lovette, I. J. (2015). New insights into New World biogeography: An integrated view from the phylogeny of blackbirds, cardinals, sparrows, tanagers, warblers, and allies. Auk, 132(2), 333–348. 10.1642/AUK-14-110.1

Barnes, R., Sahr, K., Evenden, G., Johnson, A., Warmerdam, F., Rouault, E., Song, L., & Krantz, S. (2024). dggridR: Discrete Global Grids (v. 3.1.0).

Barton, N. H., & Gale, K. S. (1993). Genetic analysis of hybrid zones. In R. G. Harrison (Ed.), Hybrid zones and the evolutionary process (pp. 13–45). Oxford University Press.

Barton, N. H., & Hewitt, G. M. (1985). Analysis of hybrid zones. Annual Review of Ecology and Systematics*. Vol.* 16, *16*, 113–148. 10.1146/annurev.es.16.110185.000553

Behr, A. A., Liu, K. Z., Liu-Fang, G., Nakka, P., & Ramachandran, S. (2016). Pong: Fast analysis and visualization of latent clusters in population genetic data. Bioinformatics, 32(18), 2817–2823. 10.1093/bioinformatics/btw327

Bouckaert, R., Vaughan, T. G., Barido-Sottani, J., Duchêne, S., Fourment, M., Gavryushkina, A., Heled, J., Jones, G., Kühnert, D., De Maio, N., Matschiner, M., Mendes, F. K., Müller, N. F., Ogilvie, H. A., Du Plessis, L., Popinga, A., Rambaut, A., Rasmussen, D., Siveroni, I., … Drummond, A. J. (2019). BEAST 2.5: An advanced software platform for Bayesian evolutionary analysis. PLoS Computational Biology, 15(4), 1–28. 10.1371/journal.pcbi.1006650

Brelsford, A., & Irwin, D. E. (2009). Incipient speciation despite little assortative mating: The yellow-rumped warbler hybrid zone. Evolution, 63(12), 3050–3060. 10.1111/j.1558-5646.2009.00777.x

Bryant, D., Bouckaert, R., Felsenstein, J., Rosenberg, N. A., & Roychoudhury, A. (2012). Inferring species trees directly from biallelic genetic markers: Bypassing gene trees in a full coalescent analysis. Molecular Biology and Evolution, 29(8), 1917–1932. 10.1093/molbev/mss086

Campagna, L., Benites, P., Lougheed, S. C., Lijtmaer, A., Giacomo, S. Di, Eaton, M. D., & Tubaro, P. L. (2012). Rapid phenotypic evolution during incipient speciation in a continental avian radiation. Proceedings of the Royal Society B: Biological Sciences, 279, 1847–1856. 10.1098/rspb.2011.2170

Campagna, L., Repenning, M., Silveira, L. F., Fontana, C. S., Tubaro, P. L., & Lovette, I. J. (2017). Repeated divergent selection on pigmentation genes in a rapid finch radiation. Science Advances, 3(5), 1–11. 10.1126/sciadv.1602404

Capblancq, T., Després, L., & Mavárez, J. (2020). Genetic, morphological and ecological variation across a sharp hybrid zone between two alpine butterfly species. Evolutionary Applications, 13(6), 1435–1450. 10.1111/eva.12925

Céspedes-Arias, L. N., Cuervo, A. M., Bonaccorso, E., Castro-Farias, M., Mendoza-Santacruz, A., Pérez-Emán, J. L., Witt, C. C., & Cadena, C. D. (2021). Extensive hybridization between two Andean warbler species with shallow divergence in mtDNA. Ornithology, 138(1), 1–28. 10.1093/ornithology/ukaa065

Chang, C. C., Chow, C. C., Tellier, L. C. A. M., Vattikuti, S., Purcell, S. M., & Lee, J. J. (2015). Second-generation PLINK: Rising to the challenge of larger and richer datasets. GigaScience, 4(1), 1–16. 10.1186/s13742-015-0047-8

Chapman, F. M. (1927). DESCRIPTIONS OF NEW BIRDS FROM NORTHWESTERN PERU AND WESTERN COLOMBIA. American Museum Novitates, 250, 1–8.

Cronemberger, Á. A., Aleixo, A., Mikkelsen, E. K., & Weir, J. T. (2020). Postzygotic isolation drives genomic speciation between highly cryptic Hypocnemis antbirds from Amazonia. In Evolution (Vol. 74, Issue 11). 10.1111/evo.14103

Cuervo, A. M. (2013). Evolutionary assembly of the Neotropical montane avifauna. Lousiana State University.

Cuervo, A. M., & Céspedes Arias, L. N. (2023). The type of *Setophaga ruficoronata* (Kaup 1851) is a hybrid: implications for the taxonomy of Myioborus warblers (Passeriformes: Parulidae). Zootaxa, 5383(4), 476–490. 10.11646/zootaxa.5383.4.3

Curson, J., David, Q., & Beadle, D. (1994). Warblers of the Americas: An Identification Guide. Houghton Mifflin Harcourt.

Danecek, P., Auton, A., Abecasis, G., Albers, C. A., Banks, E., DePristo, M. A., Handsaker, R. E., Lunter, G., Marth, G. T., Sherry, S. T., McVean, G., & Durbin, R. (2011). The variant call format and VCFtools. Bioinformatics, 27(15), 2156–2158. 10.1093/bioinformatics/btr330

DeRaad, D. A., Applewhite, E. E., Tsai, W. L. E., Terrill, R. S., Kingston, S. E., Braun, M. J., & McCormack, J. E. (2023). Hybrid zone or hybrid lineage: a genomic reevaluation of Sibley’s classic species conundrum in Pipilo towhees. Evolution, 77(3), 852–869. 10.1093/evolut/qpac068

Derryberry, E. P., Derryberry, G. E., Maley, J. M., & Brumfield, R. T. (2014). HZAR: Hybrid zone analysis using an R software package. Molecular Ecology Resources, 14(3), 652–663. 10.1111/1755-0998.12209

Edwards, S. V., Potter, S., Schmitt, C. J., Bragg, J. G., & Moritz, C. (2016). Reticulation, divergence, and the phylogeography-phylogenetics continuum. Proceedings of the National Academy of Sciences of the United States of America, 113(29), 8025–8032. 10.1073/pnas.1601066113

Fitak, R. R. (2021). OptM: Estimating the optimal number of migration edges on population trees using Treemix. Biology Methods and Protocols, 6(1), 1–6. 10.1093/biomethods/bpab017

Flantua, S. G. A., Hooghiemstra, H., Boxel, J. H. Van, Cabrera, M., González Carranza, Z., & González-Arango, C. (2014). Connectivity dynamics since the Last Glacial Maximum in the northern Andes; a pollen-driven framework to assess potential migration. In W. D. Stevens, O. M. Montiel, & P. Raven (Eds.), Paleobotany and Biogeography: A Festschrift for Alan Graham in His 80th Year (pp. 98–123). Missouri Botanical Garden Press.

Friis, G., Aleixandre, P., Rodríguez-Estrella, R., Navarro-Sigüenza, A. G., & Milá, B. (2016). Rapid postglacial diversification and long-term stasis within the songbird genus Junco: phylogeographic and phylogenomic evidence. Molecular Ecology, 25(24), 6175–6195. 10.1111/mec.13911

Gay, L., Crochet, P. A., Bell, D. A., & Lenormand, T. (2008). Comparing clines on molecular and phenotypic traits in hybrid zones: A window on tension zone models. Evolution, 62(11), 2789–2806. 10.1111/j.1558-5646.2008.00491.x

Gompert, Z., DeRaad, D. A., & Buerkle, C. A. (2024). A next generation of hierarchical Bayesian analyses of hybrid zones enables direct quantification of variation in introgression in R. BioRxiv, 2003–2024. 10.1002/ece3.70548

Graham, C. H., Silva, N., & Velásquez-Tibatá, J. (2010). Evaluating the potential causes of range limits of birds of the Colombian Andes. Journal of Biogeography, 37(10), 1863–1875. 10.1111/j.1365-2699.2010.02356.x

Graves, G. R. (1988). Linearity of geographic range and its possible effect on the population structure of Andean birds. The Auk, 105(1), 47–52.

Gutiérrez-Pinto, N., Cuervo, A. M., Miranda, J., Pérez-Emán, J. L., Brumfield, R. T., & Cadena, C. D. (2012). Non-monophyly and deep genetic differentiation across low-elevation barriers in a Neotropical montane bird (*Basileuterus tristriatus*; Aves: Parulidae). Molecular Phylogenetics and Evolution, 64(1), 156–165. 10.1016/j.ympev.2012.03.011

Gutiérrez-Pinto, N., Cuervo, A. M., Miranda, J., Pérez-Emán, J. L., Brumfield, R. T., Cadena, C. D., Gutiérrez-Pinto, N., Cuervo, A. M., Miranda, J., & Pérez-Emán, J. L. (2017). Non-monophyly and deep genetic differentiation across low-elevation barriers in a Neotropical montane bird … Molecular Phylogenetics and Evolution, 64(August), 156–165. 10.1016/j.ympev.2012.03.011

Harrison, R. G. (1986). Pattern and process in a narrow hybrid zone. Heredity, 56(3), 337–349. 10.1038/hdy.1986.55

Hazzi, N. A., Moreno, J. S., Ortiz-Movliav, C., & Palacio, R. D. (2018). Biogeographic regions and events of isolation and diversification of the endemic biota of the tropical Andes. Proceedings of the National Academy of Sciences of the United States of America, 115(31), 7985–7990. 10.1073/pnas.1803908115

Irwin, D. E., Rubtsov, A. S., & Panov, E. N. (2009). Mitochondrial introgression and replacement between yellowhammers (*Emberiza citrinella*) and pine buntings (*Emberiza leucocephalos*) (Aves: Passeriformes). Biological Journal of the Linnean Society, 98, 422–438.

Jenkins, T. L. (2024). mapmixture: An R package and web app for spatial visualisation of admixture and population structure. Molecular Ecology Resources, 24(4), 1–6. 10.1111/1755-0998.13943

Kearns, A. M., Restani, M., Szabo, I., Schrøder-Nielsen, A., Kim, J. A., Richardson, H. M., Marzluff, J. M., Fleischer, R. C., Johnsen, A., & Omland, K. E. (2018). Genomic evidence of speciation reversal in ravens. Nature Communications, 9(1). 10.1038/s41467-018-03294-w

Li, H., & Durbin, R. (2009). Fast and accurate short read alignment with Burrows-Wheeler transform. Bioinformatics, 25(14), 1754–1760. 10.1093/bioinformatics/btp324

Lim, H. C., Bennett, K. F. P., Justyn, N. M., Powers, M. J., Long, K. M., Kingston, S. E., Lindsay, W. R., Pease, J. B., Fuxjager, M. J., Bolton, P. E., Balakrishnan, C. N., Day, L. B., Parsons, T. J., Brawn, J. D., Hill, G. E., & Braun, M. J. (2024). Sequential introgression of a carotenoid processing gene underlies sexual ornament diversity in a genus of manakins. Science Advances, 10(47). 10.1126/sciadv.adn8339

Linck, E., & Battey, C. J. (2019). Minor allele frequency thresholds strongly affect population structure inference with genomic data sets. Molecular Ecology Resources, 19(3), 639–647. 10.1111/1755-0998.12995

Liu, S., Zhang, L., Sang, Y., Lai, Q., Zhang, X., Jia, C., Long, Z., Wu, J., Ma, T., Mao, K., Street, N. R., Ingvarsson, P. K., Liu, J., & Wang, J. (2022). Demographic History and Natural Selection Shape Patterns of Deleterious Mutation Load and Barriers to Introgression across Populus Genome. Molecular Biology and Evolution, 39(2), 1–16. 10.1093/molbev/msac008

Long, K. M., Rivera-Colón, A. G., Bennett, K. F. P., Catchen, J. M., Braun, M. J., & Brawn, J. D. (2024). Ongoing introgression of a secondary sexual plumage trait in a stable avian hybrid zone. Evolution, 78(9), 1539–1553. 10.1093/evolut/qpae076

Lopez, K. A., McDiarmid, C. S., Griffith, S. C., Lovette, I. J., & Hooper, D. M. (2021). Evaluating evidence of mitonuclear incompatibilities with the sex chromosomes in an avian hybrid zone. Evolution, 75(6), 1395–1414. 10.1111/evo.14243

Lovette, I. J., Pérez-Emán, J. L., Sullivan, J. P., Banks, R. C., Fiorentino, I., Córdoba-Córdoba, S., Echeverry-Galvis, M., Barker, F. K., Burns, K. J., Klicka, J., Lanyon, S. M., & Bermingham, E. (2010). A comprehensive multilocus phylogeny for the wood-warblers and a revised classification of the Parulidae (Aves). Molecular Phylogenetics and Evolution, 57(2), 753–770. 10.1016/j.ympev.2010.07.018

Malinsky, M., Trucchi, E., Lawson, D. J., & Falush, D. (2018). RADpainter and fineRADstructure: Population Inference from RADseq Data. Molecular Biology and Evolution, 35(5), 1284–1290. 10.1093/molbev/msy023

Manichaikul, A., Mychaleckyj, J. C., Rich, S. S., Daly, K., Sale, M., & Chen, W. M. (2010). Robust relationship inference in genome-wide association studies. Bioinformatics, 26(22), 2867–2873. 10.1093/bioinformatics/btq559

Marchetti, K. (1992). Dark habitats and bright birds illustrate the role of the environment in species divergence. Nature, 362, 149–152.

Marcus, J., Ha, W., Barber, R. F., & Novembre, J. (2021). Fast and flexible estimation of effective migration surfaces. ELife, 10, 1–46. 10.7554/eLife.61927

Márquez, R., Linderoth, T. P., Mejía-Vargas, D., Nielsen, R., Amézquita, A., & Kronforst, M. R. (2020). Divergence, gene flow, and the origin of leapfrog geographic distributions: The history of colour pattern variation in Phyllobates poison-dart frogs (preprint). In BioRxiv. 10.1111/mec.15598

Martínez-Gómez, S. C., Lara, C. E., Remsen, J. V., Brumfield, R. T., & Cuervo, A. M. (2023). Unmasking hidden genetic, vocal, and size variation in the Masked Flowerpiercer along the Andes supports two species separated by Northern Peruvian Low. Ornithology, 140(4), 1–14. 10.1093/ornithology/ukad028

Mayr, E. (1942). Systematics and the origin of species from the viewpoint of a zoologist. Columbia University Press.

Meirmans, P. G. (2012). The trouble with isolation by distance. Molecular Ecology, 21(12), 2839–2846. 10.1111/j.1365-294X.2012.05578.x

Molloy, E. K., Durvasula, A., & Sankararaman, S. (2021). Advancing admixture graph estimation via maximum likelihood network orientation. Bioinformatics, 37, I142–I150. 10.1093/bioinformatics/btab267

Mumme, R. L. (2015). Demography of Slate-throated Redstarts (*Myioborus miniatus*): a non-migratory Neotropical warbler. Journal of Field Ornithology, 86(2), 89–102. 10.1111/jofo.12093

Natola, L., Curtis, A., Hudon, J., & Burg, T. M. (2021). Introgression between Sphyrapicus nuchalis and S. varius sapsuckers in a hybrid zone in west-central Alberta. Journal of Avian Biology, 52(8), 1–12. 10.1111/jav.02717

Nosil, P. (2008). Ernst Mayr and the integration of geographic and ecological factors in speciation. Biological Journal of the Linnean Society, 95(1), 26–46. 10.1111/j.1095-8312.2008.01091.x

Nychka, D., Furrer, R., Paige, J., Sain, S., & Nychka, M. D. (2025). Package ‘fields.’ 10.5065/D6W957CT>.

Oksanen, J., Kindt, R., Legendre, P., O’Hara, B., Stevens, M. H. H., Oksanen, M. J., & Suggests, M. A. S. S. (2007). *“*The vegan package*.”* Community ecology package.

Paris, J. R., Stevens, J. R., & Catchen, J. M. (2017). Lost in parameter space: a road map for stacks. Methods in Ecology and Evolution, 8(10), 1360–1373. 10.1111/2041-210X.12775

Pérez-Emán, J. L. (2005). Molecular phylogenetics and biogeography of the Neotropical redstarts (*Myioborus*; Aves, Parulinae). Molecular Phylogenetics and Evolution, 37(2), 511–528. 10.1016/j.ympev.2005.04.013

Peterson, B. K., Weber, J. N., Kay, E. H., Fisher, H. S., & Hoekstra, H. E. (2012). Double digest RADseq: An inexpensive method for de novo SNP discovery and genotyping in model and non-model species. PLoS ONE, 7(5). 10.1371/journal.pone.0037135

Pickrell, J. K., & Pritchard, J. K. (2012). Inference of Population Splits and Mixtures from Genome-Wide Allele Frequency Data. PLoS Genetics, 8(11). 10.1371/journal.pgen.1002967

Poelstra, J. W., Vijay, N., Bossu, C. M., Lantz, H., Ryll, B., Müller, I., Baglione, V., Unneberg, P., Wikelski, M., Grabherr, M. G., & Wolf, J. B. W. (2014). The genomic landscape underlying phenotypic integrity in the face of gene flow in crows. Science, 344(6190), 1410–1414. 10.1126/science.1253226

Pool, J. E., & Nielsen, R. (2009). Inference of historical changes in migration rate from the lengths of migrant tracts. Genetics, 181(2), 711–719. 10.1534/genetics.108.098095

Pozo-Zamora, G. M., Krabbe, N., Mena-Valenzuela, P., Nilsson, J., & Brito, J. (2022). Birds of the Cordillera del Kutukú, Morona Santiago, Southeastern Ecuador. Revista Peruana de Biologia, 29(1). 10.15381/RPB.V29I1.20667

Price, T. (2008). Speciation in Birds. Robert and Co.

Prieto-Torres, D. A., Cuervo, A. M., & Bonaccorso, E. (2018). On geographic barriers and Pleistocene glaciations : Tracing the diversification of the Russet-crowned Warbler (*Myiothlypis coronata*) along the Andes. Plos One, 13(3), e0191598.

Rahbek, C., Borregaard, M. K., Colwell, R. K., Dalsgaard, B., Holt, B. G., Morueta-Holme, N., Nogues-Bravo, D., Whittaker, R. J., & Fjeldså, J. (2019). Humboldt’s enigma: What causes global patterns of mountain biodiversity? Science, 365(6458), 1108–1113. 10.1126/science.aax0149

Rambaut, A., Suchard, M. A., Xie, D., & Drummond, A. J. (2014). Tracer v1.6.

Ramírez-Barahona, S., & Eguiarte, L. E. (2013). The role of glacial cycles in promoting genetic diversity in the Neotropics : the case of cloud forests during the Last Glacial Maximum. Ecology and Evolution, 3(3), 725–738. 10.1002/ece3.483

Remsen, J. V. (1984). High Incidence of “Leapfrog” Pattern of Geographic variation in Andean Birds: Implications for the speciation process. Science, 224, 171–173.

Rémy, A., Grégoire, A., Perret, P., & Doutrelant, C. (2010). Mediating male-male interactions: The role of the UV blue crest coloration in blue tits. Behavioral Ecology and Sociobiology, 64(11), 1839–1847. 10.1007/s00265-010-0995-z

Rivera-Colón, A. G., & Catchen, J. (2022). Population Genomics Analysis with RAD, Reprised: Stacks 2 (Chapter 7). In C. Verde & D. Giordano (Eds.), Marine Genomics: Methods and Protocols. Springer Nature-Humana Press.

Robbins, M. B., Ridgely, R. S., & Schulenberg, T. S. (1987). The avifauna of the Cordillera de Cutucut, Ecuador, with Comparisons to other Andean Localities. Proceedings of The Academy of Natural Sciences of Philadelphia, 139(1), 243–259.

Rochette, N. C., Rivera-Colón, A. G., & Catchen, J. M. (2019). Stacks 2: Analytical methods for paired-end sequencing improve RADseq-based population genomics. Molecular Ecology, 28(21), 4737–4754. 10.1111/mec.15253

Sambrook, J. (1987). Molecular cloning: a laboratory manual.

Sanín, M. J., Cardona, A., Céspedes-Arias, L. N., González-Arango, C., Pardo, N., & Cadena, C. D. (2024). Volcanoes, evolving landscapes, and biodiversity in Neotropical mountains. Frontiers of Biogeography, 16(1), 1–14. 10.21425/F5FBG61882

Scordato, E. S. C., Wilkins, M. R., Semenov, G., Rubtsov, A. S., Kane, N. C., & Safran, R. J. (2017). Genomic variation across two barn swallow hybrid zones reveals traits associated with divergence in sympatry and allopatry. Molecular Ecology, 26(20), 5676–5691. 10.1111/mec.14276

Scott, G. W., & Deag, J. M. (1998). Blue Tit (*Parus caeruleus*) agonistic displays : A reappraisal. Behaviour, 135(6), 665–691.

Seeholzer, G. F., & Brumfield, R. T. (2018). Isolation by distance, not incipient ecological speciation, explains genetic differentiation in an Andean songbird (Aves: Furnariidae: Cranioleuca antisiensis, Line-cheeked Spinetail) despite near threefold body size change across an environmental gradien. Molecular Ecology, 27(1), 279–296. 10.1111/mec.14429

Semenov, G. A., Linck, E., Enbody, E. D., Harris, R. B., Khaydarov, D. R., Alström, P., Andersson, L., & Taylor, S. A. (2021). Asymmetric introgression reveals the genetic architecture of a plumage trait. Nature Communications, 12(1), 1–9. 10.1038/s41467-021-21340-y

Sonne, J., Dalsgaard, B., Borregaard, M. K., Kennedy, J., Fjeldså, J., & Rahbek, C. (2022). Biodiversity cradles and museums segregating within hotspots of endemism. Proceedings of the Royal Society B: Biological Sciences, 289(1981). 10.1098/rspb.2022.1102

Stein, A. C., & Uy, J. A. C. (2006). Plumage brightness predicts male mating success in the lekking golden-collared manakin, Manacus vitellinus. Behavioral Ecology, 17(1), 41–47. 10.1093/beheco/ari095

Stoltz, M., Baeumer, B., Bouckaert, R., Fox, C., Hiscott, G., & Bryant, D. (2021). Bayesian inference of species trees using diffusion models. Systematic Biology, 70(1), 145–161. 10.1093/sysbio/syaa051

Stryjewski, K. F., & Sorenson, M. D. (2017). Mosaic genome evolution in a recent and rapid avian radiation. Nature Ecology and Evolution, 1(12), 1912–1922. 10.1038/s41559-017-0364-7

Thrasher, D. J., Butcher, B. G., Campagna, L., Webster, M. S., & Lovette, I. J. (2018). Double-digest RAD sequencing outperforms microsatellite loci at assigning paternity and estimating relatedness: A proof of concept in a highly promiscuous bird. Molecular Ecology Resources, 18(5), 953–965. 10.1111/1755-0998.12771

Toews, D. P. L., & Brelsford, A. (2012). The biogeography of mitochondrial and nuclear discordance in animals. Molecular Ecology, 21, 3907–3930. 10.1111/j.1365-294X.2012.05664.x

Toews, D. P. L., Campagna, L., Taylor, S. A., Balakrishnan, C. N., Baldassere, D. T., Deane-Coe, P. E., Harvey, M. G., Hooper, D. M., Irwin, D. E., Judy, C. D., Mason, N. A., McCormack, J. E., McCracken, K. G., Oliveros, C. H., Safran, R. J., Scordato, E. S. C., Stryjewski, K. F., Tigano, A., Uy, J. A. C., & Winger, B. (2016). Genomic approaches to understanding the early stages of population divergence and speciation in birds. The Auk, 133, 13–30. 10.1642/AUK-15-51.1

Toews, D. P. L., Taylor, S. A., Vallender, R., Brelsford, A., Butcher, B., Messer, P. W., & Lovette, I. J. (2016). Plumage genes and little else distinguish the genomes of hybridizing warblers. Current Biology, 26(17), 2313–2318. 10.1016/j.cub.2016.06.034

Uy, J. A. C., Irwin, D. E., & Webster, M. S. (2018). Behavioral isolation and incipient speciation in birds. Annual Review of Ecology, Evolution, and Systematics, 49, 1–24. 10.1146/annurev-ecolsys-110617-062646

Vos, M., & Velicer, G. J. (2008). Isolation by Distance in the Spore-Forming Soil Bacterium Myxococcus xanthus. Current Biology, 18(5), 386–391. 10.1016/j.cub.2008.02.050

Vuilleumier, F. (1969). Pleistocene speciation of birds living in the high Andes. Nature, 223, 1179–1180.

Walsh, J., Lovette, I. J., Winder, V., Elphick, C. S., Olsen, B. J., Shriver, G., & Kovach, A. I. (2017). Subspecies delineation amid phenotypic, geographic and genetic discordance in a songbird. Molecular Ecology, 26(5), 1242–1255. 10.1111/mec.14010

Weir, B., & Cockerham, C. (1984). Estimating F-Statistics for the Analysis of Population Structure. Evolution, 38(6), 1358–1370.

Wiens, B. J., & Colella, J. P. (2025). triangulaR: an R package for identifying AIMs and building triangle plots using SNP data from hybrid zones. Heredity, 134, 251–262. 10.1038/s41437-025-00760-2

Winger, B. M. (2017). Consequences of divergence and introgression for speciation in Andean cloud forest birds. Evolution, 71(7), 1815–1831. 10.1111/evo.13251

Winger, B. M., & Bates, J. M. (2015). The tempo of trait divergence in geographic isolation: Avian speciation across the Marañon Valley of Peru. Evolution, 69(3), 772–787. 10.1111/evo.12607

World, H. of the B. of the, & International, B. (2019). Handbook of the Birds of the World and BirdLife International digital checklist of the birds of the world. Version 4.

Yi, X., & Latch, E. K. (2022). Nonrandom missing data can bias Principal Component Analysis inference of population genetic structure. Molecular Ecology Resources, 22(2), 602–611. 10.1111/1755-0998.13498

Zamudio, K. R., Bell, R. C., & Mason, N. A. (2016). Phenotypes in phylogeography: Species’ traits, environmental variation, and vertebrate diversification. Proceedings of the National Academy of Sciences of the United States of America, 113(29), 8041–8048. 10.1073/pnas.1602237113

Zheng, X. (2013). Package ‘SNPRelate’: A High-performance computing toolset for relatedness and principal component analysis of SNP data.

Zimmer, J. T. (1949). Studies of Peruvian birds. No. 54, The families Catamblyrhynchidae and Parulidae. American Museum Novitates, 1428, 1–59.

