## Supplementary Material for "History of divergence and gene flow shaping geographic variation in Andean warblers (*Myioborus*)"

### Supplementary information- History of divergence and gene flow shaping geographic variation in color in Andean warbles

#### Supplementary information- Methods

##### DNA extraction and library preparation (full)

We extracted genomic DNA from pectoral muscle (n=232) and blood samples (n=3) employing a modified phenol-chlorophorm protocol (Sambrook 1987), using phase-lock tubes. Each DNA extraction was subsequently purified using homemade magnetic nucleic acid extraction (MagNA) made with Sera-Mag Magnetic Speed-beads. To obtain reduced representation genomic data, we used double-digest restriction site-associated DNA sequencing (ddRAD) following Peterson et al. (2012), and using modifications from Thrasher et al. (2018). Before preparing the genomic libraries, we measured DNA concentrations using a Qubit fluorometer (Life Technologies, Carlsbad, California, USA). For library preparation, we used approximately 500 ng of DNA per extraction, which had concentrations ranging from 6.86–55 ng/μL. Extractions of four individuals for which we had concerns about quality were included twice in the libraries with a different barcode (i.e., there were a total of 239 samples included, corresponding to 235 individuals). After demultiplexing, the reads corresponding to the same individual were concatenated into the same file (see below).

We prepared libraries as follows. First, DNA was digested by incubating the samples with the enzymes SbfI (8 base pair [bp] recognition site) and MspI (4 bp recognition site; New England BioLabs, Ipswich, Massachusetts, USA). Simultaneously, the P1 and P2 adaptors were ligated using T4 DNA Ligase (New England BioLabs). After digestion and ligation, samples with unique P1 barcodes were pooled together in 12 index groups of 20 individuals, each identified with a unique P2 adapter. After pooling, we purified the DNA in each index group using homemade MagNA (1.5X volumes). Then DNA fragments between 450 and 600 bp were selected using Blue Pippin, by the Cornell University Biotechnology Resource Center (BRC). After the size selection step, we incorporated the index group and Illumina sequencing adapters by conducting a PCR (as in Thrasher et al. 2018). We then purified the reactions using MagNA, and pooled equimolarly to obtain a single library for Illumina sequencing.

#### Bioinformatics pipeline

**Optimization of *de novo* assembly parameters (full)-** Before assembling the reads, we conducted an optimization of the parameters *M* and *n* used by *denovo_map.pl* (Rochette et al. 2019). These parameters determine the number of mismatches allowed between stacks to merge into putative loci, and between loci when building the catalog, respectively (https://catchenlab.life.illinois.edu/stacks/comp-v1/denovo_map.php ). Adequate values of these parameters are highly dependent on the dataset's characteristics, such as the genetic diversity and level of divergence among populations, and therefore it is necessary to perform this optimization step for *de novo* assemblies (Paris et al. 2017; Rivera-Colón and Catchen 2022). For this reason, we also conducted three separate optimizations and then, three separate full assemblies: one including representatives from the outgroups, one that only included individuals within the *ornatus-melanocephalus* complex, and one including only samples relevant for the hybrid zone analyses (*chrysops*, individuals in the hybrid zone, *bairdi,* and *griseonuchus*, Figure 1).
We subset the dataset retaining a small number of representative individuals for the three optimization runs (Rivera-Colón and Catchen 2022). Specifically, we included at least one sample per taxon within the complex, and only those that had an average number of reads after

demultiplexing (+/- 0.25 of the standard deviation). These led to population maps with 43 (the full, i.e. including outgroups), 40 (ingroup only), and 32 (hybrid zone only) assemblies. In these three optimization population maps, all of the individuals are grouped into one "population" (Rivera-Colón and Catchen 2022). To optimize *M* and *n* we ran *denovo-map.pl* using 36 combinations of these parameters following recommendations by Paris *et al* (2017) and Rivera-Colón & Catchen (2022): we varied *M* from 1 to 12, and then each value of M was run together with three *n* values (*n=M*, *n=M+1* and *n=M-1*). The parameter *m* was kept constant at the default value that has proven to work well in a diversity of biological scenarios (Paris et al. 2017). In each case we chose the *M/n* combination that maximized the number of polymorphic loci that were shared by at least 80% of the individuals (Paris et al. 2017; Rivera-Colón and Catchen 2022). Ultimately, we chose M=4/n=5 for the full assembly, M=3/n=4 for the only ingroup assembly, and M=3/n=4 for the hybrid zone assembly (Supplementary Table 2-denovo_optimization_results.xls).

**SNP datasets (full)-** We obtained different SNP datasets to be used in downstream analyses using the *populations* program from STACKS 2.66 (Rochette et al. 2019). For some analyses, as specified below, we did further filtering of SNPs using *vcftools* (Danecek et al. 2011). For the ingroup assembly, we obtained two different SNP datasets: one including one SNP per locus, and one that included all variant and non-variant sites per locus (i.e. full haplotypes only for fineRADStructure, see below). In both cases we applied the following filters: *--min-samples-overall* 0.8 (locus needs to be present in 80% of all individuals to be retained), *--min-mac* 5 (minimum allele count of 5), and *--max-obs-het* 0.7 (maximum observed heterozygosity of 0.7). For the full haplotype dataset, we used the same parameters and included the *--filter-haplotype-*

*wise* flag. We decided to apply a minimum allele count (*--min-mac* 5) filter to exclude SNPs that may be artifacts of genotyping errors, but kept a relatively low value to avoid excluding all low frequency SNPs which also contain relevant information for population genetic analyses (Linck and Battey 2019).

The assembly including outgroups was only used for phylogenetic (*raxml-ng*, and *snapper*) and admixture graph inference (see below). We obtained slightly different SNP datasets for each analysis, but for all we applied the following filters: *--min-samples-per-pop* 0.5, *--min-populations* {total number of populations in population map, either 10 or 8}, *--min-mac* 2, and *--max-obs-het* 0.7. In this case we lowered the *--min-mac* 2 value because outgroups are represented only by 2 or 3 individuals, and these likely introduce very low frequency alleles. Additional parameters used during filtering are specified below in the corresponding analyses. The "hybrid zone assembly" was used for analyses aimed at describing genetic variation across the *bairdi-chrysops* hybrid zone and adjacent areas. First, informed by our whole complex results (e.g. fineRADStructure, see results), we decided to conduct PCA and admixture analyses for this subset of individuals to further describe genetic structure in southern Ecuador, which is relevant for the patterns of genetic variation across the *bairdi x chrysops* hybrid zone. For this set of analyses, we obtained a dataset including one SNP per locus applying the following filtering: -*min-samples-overall* 0.8, *--min-mac* 5 and *--max-obs-het* 0.7. For hybrid zone-specific analyses (i.e. cline and triangulaR analyses, see below) we obtained a SNP dataset excluding all *griseonuchus* (from northern Peru) as well as individuals from southern Ecuador that genetically cluster with *griseonuchus* (see results). For these hybrid zone-specific analyses, we obtained SNP datasets both including one SNP and all SNPs per locus for different analyses. We used the same SNP filtering scheme as for the SNP dataset including *griseonuchus*.

Based on the assembly for the ingroup only, and after filtering, we obtained 5,054 loci, composed of 73,786 sites, including 22,132 variant sites for the full haplotype dataset. For the ingroup (one SNP per locus) dataset, we obtained the same number of loci and kept 4,800 variant sites. The number of SNPs for the other two assemblies (i.e., including outgroups and for the hybrid zone only), as well as for different filtering employed for different analyses is fully detailed in the Supplementary Table 2

#### References


Danecek, P., A. Auton, G. Abecasis, C. A. Albers, E. Banks, M. A. DePristo, R. E. Handsaker, G. Lunter, G. T. Marth, S. T. Sherry, G. McVean, and R. Durbin. 2011. The variant call format and VCFtools. Bioinformatics 27:2156–2158.

Linck, E., and C. J. Battey. 2019. Minor allele frequency thresholds strongly affect population structure inference with genomic data sets. Mol. Ecol. Resour. 19:639–647.

Paris, J. R., J. R. Stevens, and J. M. Catchen. 2017. Lost in parameter space: a road map for stacks. Methods Ecol. Evol. 8:1360–1373.

Peterson, B. K., J. N. Weber, E. H. Kay, H. S. Fisher, and H. E. Hoekstra. 2012. Double digest RADseq: An inexpensive method for de novo SNP discovery and genotyping in model and non-model species. PLoS One 7.

Rivera-Colón, A. G., and J. Catchen. 2022. Population Genomics Analysis with RAD, Reprised: Stacks 2 (Chapter 7). P. *in* C. Verde and D. Giordano, eds. Marine Genomics: Methods and Protocols. Springer Nature- Humana Press.

Rochette, N. C., A. G. Rivera-Colón, and J. M. Catchen. 2019. Stacks 2: Analytical methods for paired-end sequencing improve RADseq-based population genomics. Mol. Ecol. 28:4737–

4754.

Sambrook, J. 1987. Molecular cloning: a laboratory manual. New York.

Thrasher, D. J., B. G. Butcher, L. Campagna, M. S. Webster, and I. J. Lovette. 2018. Double-digest RAD sequencing outperforms microsatellite loci at assigning paternity and estimating relatedness: A proof of concept in a highly promiscuous bird. Mol. Ecol. Resour. 18:953–965.

### Supplementary Figures

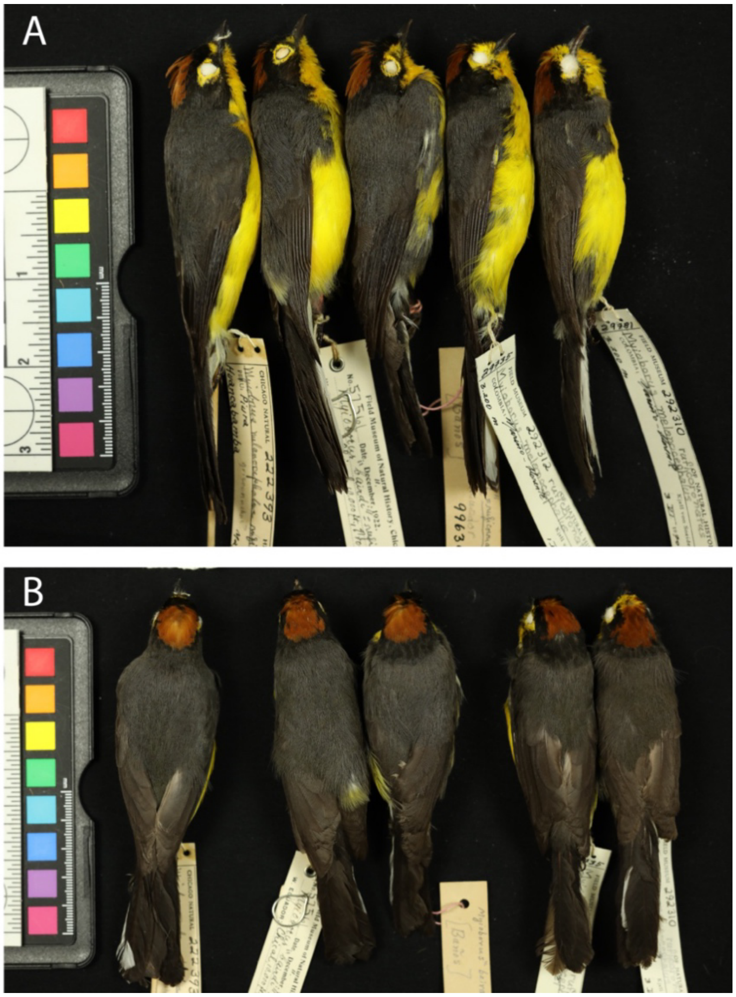


**Figure S1.** Sample of plumage variation from northern Peru (Piura) to southern Colombia (Nariño). The first specimen, from left to right, corresponding clearly to the *griseonuchus* group, shows a wide black band below the yellow spectacles that extend down to the cheeks, "broken" yellow spectacles and lacks a black band in the nape (i.e. the rufous crown is in direct contact with the grayish back color). The two specimens in the middle, collected in the known distribution of the *bairdi* group, also show a complete black band below the yellow spectacles, but vary on the extent that these extend down to the cheek (with the specimen to the left being more *griseonuchus*-like). In both cases the yellow spectacles are not broken. The two specimens also differ in the extent of the black nape, which is also one of the described differences between these plumage forms. The specimen closer to the left shows a very narrow black nape band, somehow resembling the *griseonuchus* specimen. The last two specimens, from left to right, were collected in the known *chrysops x bairdi* hybrid zone (see more in Figure 7 about plumage variation in the hybrid zone. The specimens shown from the left to the right: FMNH 222393 (Huancabamba, dept. Piura, Peru); FMNH 57561 (Chical , Prov. Cañar, Ecuador); FMNH 9963 (Baños, prov. Tungurahua, Ecuador); FMNH 292312 and 29310 (Llorente, depto. Nariño, Colombia).

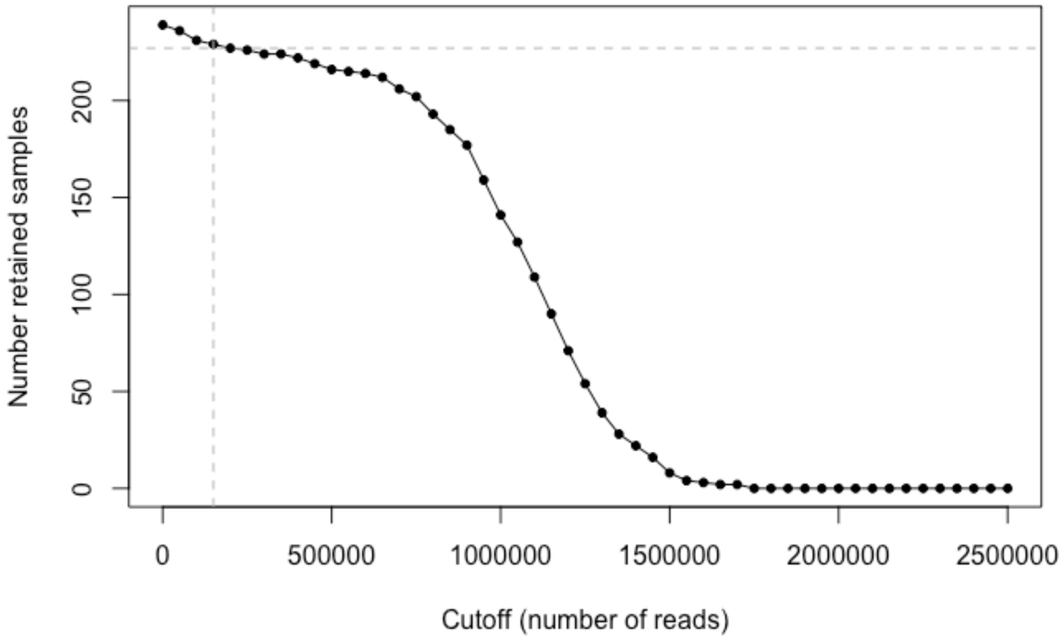


**Figure S2.** Number of retained samples based on different cutoffs of number of reads. We chose a cut off of 200,000 reads, shown in the dotted vertical line.

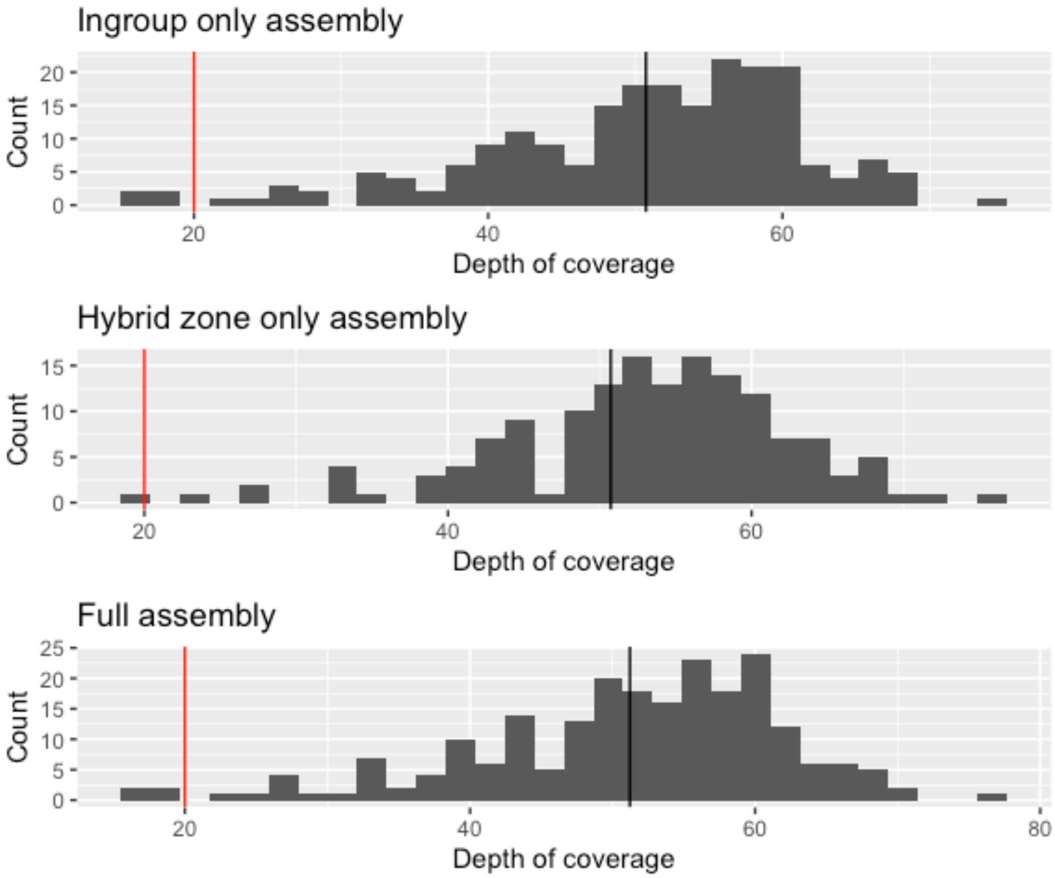


**Figure S3.** Distribution of coverage for the three assemblies. The red vertical line shows the lower threshold for coverage: all individual below this threshold were discarded from downstream analyses. The black vertical line shows the mean of depth of coverage.

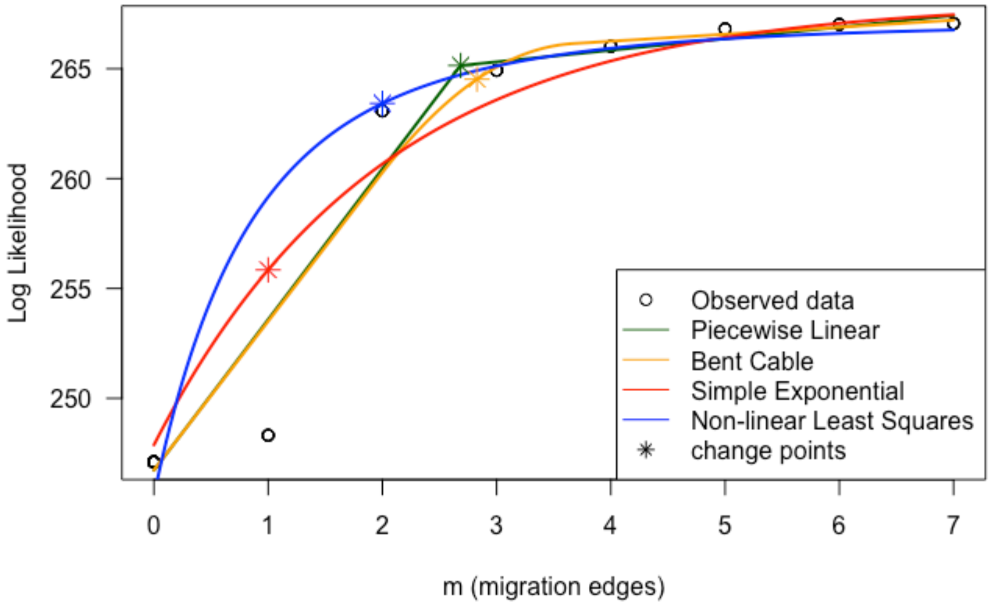


**Figure S4.** Log-likelihood across different values of migration edges (*OrientAGraph*). The asterisk shows inflection points that can be interpreted as the optimal m values. We show the results for m=1 and m=3, in Figure 2.

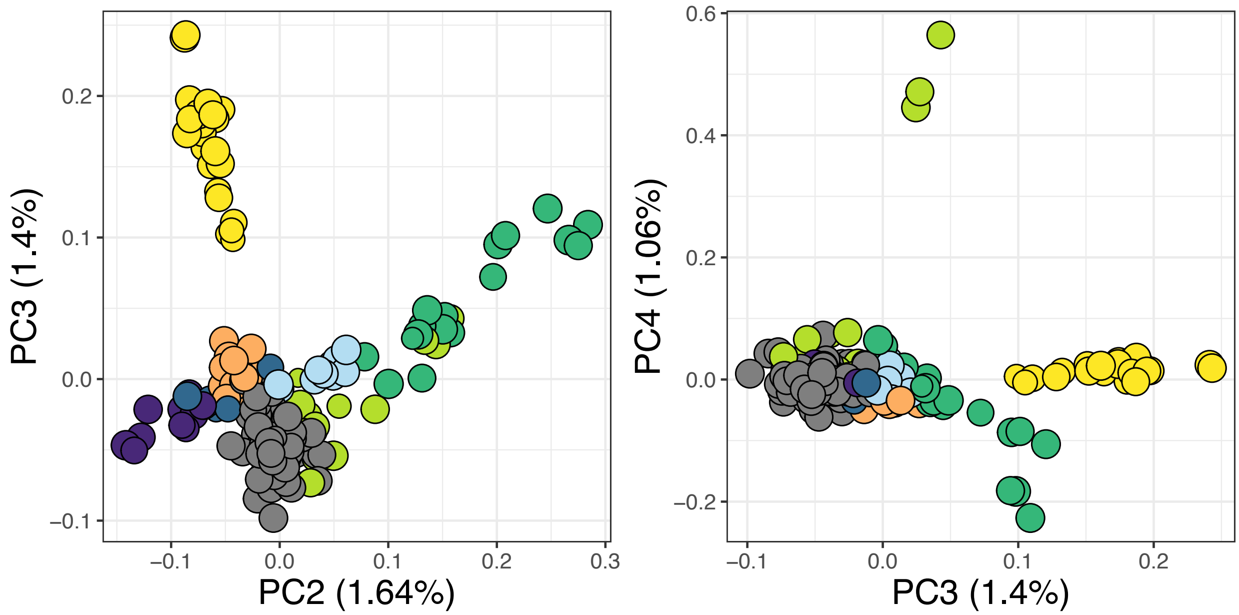


**Figure S5.** Distribution of individuals in the PC2/PC3 and PC3/PC4 spaces, showing clustering by plumage group which is encoded by color (as in main figures.)

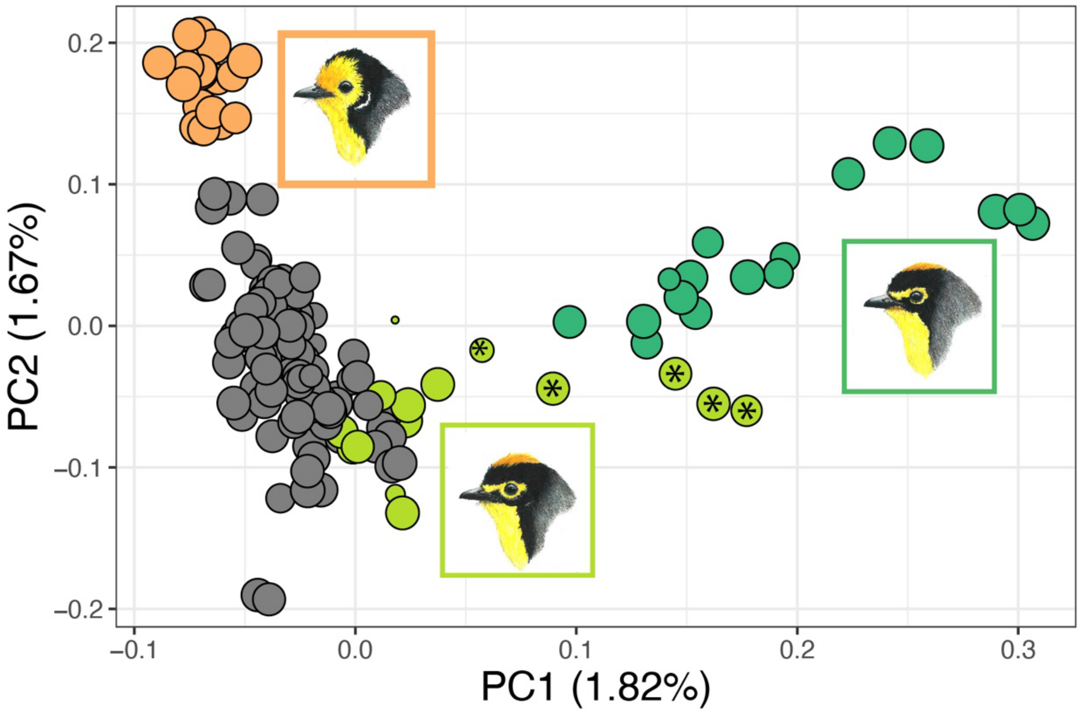


**Figure S6.** Genetic structure in the hybrid zone, parental populations and griseonuchus, as revealed by a PCA. The asterisks denote individuals from Loja (Ecuador) and outlining ridges in Morona-Santiago that were initially classified as *bairdi* but genetically cluster with *griseonuchus* in other analyses (e.g. fineRADStructure).

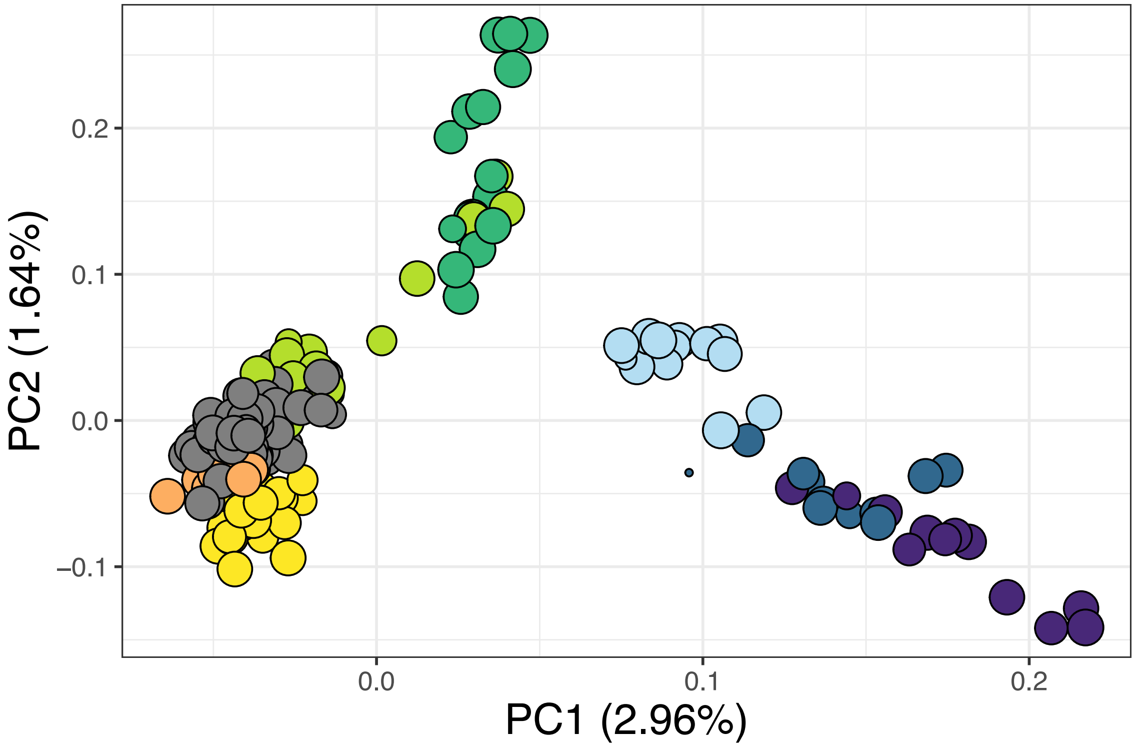


**Figure S7.** Distribution of individuals across the PC1/PC2 space, using a SNP dataset that excludes SNPs with high levels of missing data. Plumage groups are denoted by color.

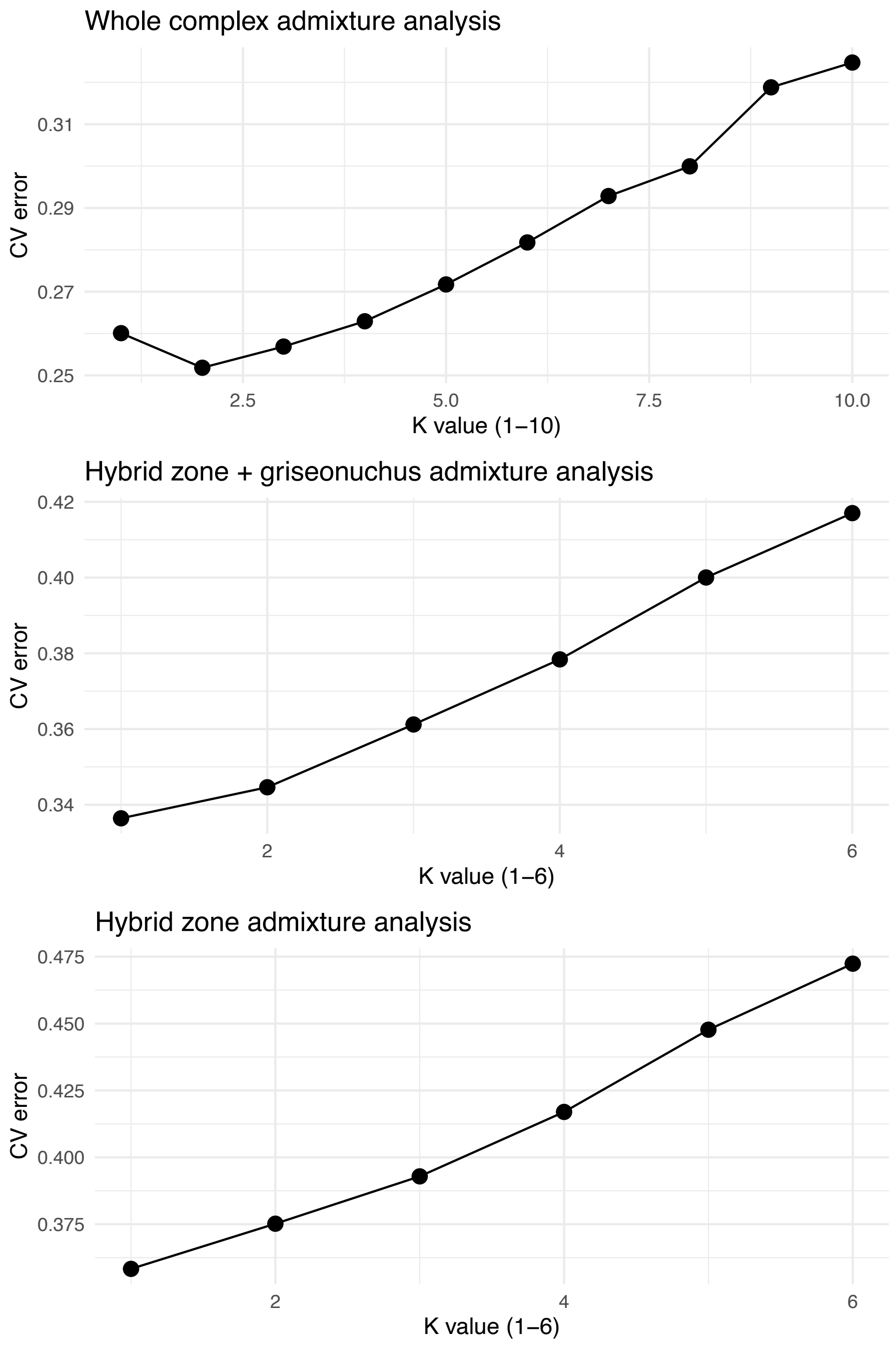


**Figure S8.** Cross-validation procedure results for admixture analyses.

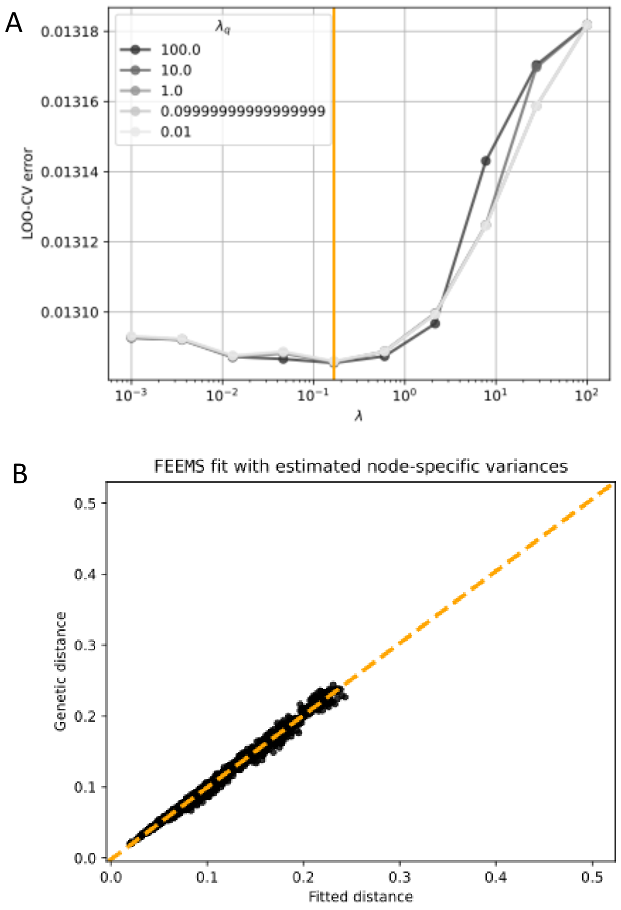


**Figure S9.** FEEMS model parametrization and fit. **A)** Cross-validation (CV) error plot used to select the optimal graph penalty parameter, with the chosen value indicated. **B)**Observed genetic distances plotted against model-fitted effective distances, showing the goodness-of-fit of the inferred migration surface.

### Supplementary Tables (captions)

**Table S1.** Metadata for each of the sampled individuals. Columns include:

- **RAD_ID, Lab_ID, Tissue_ID, Skin_ID** – Sample identifiers across datasets and tissue types.
- **Sex** – Recorded sex of individual.
- **Source** – Museum or collection source.
- **Species, Species_Zootaxa** – Species identification and classification. The "Species_Zootaxa" column corresponds to a classification based on reccomendations by Cuervo & Céspedes 2023.
- **Group_geography** – Geographic group assignment (i.e. subspecies or hybrid zone bird)
- **populations_1, populations_2** – Population designations for different analyses.
- **Country, Department, Locality** – Geographic origin of sample.
- **Locality_Construct, Locality_HZanalyses** – Locality groupings used for population and hybrid zone analyses.
- **Latitude, Longitude, coord_general_check** – Geographic coordinates and coordinate validation.

- **ddRAD_kept_postDemult, included_finalAnalyses** – Inclusion status in ddRAD pipeline and final analyses.
- **Group_plumage, Base_plumage_record** – Plumage-based group and reference record.
- **HI** – Plumage hybrid index
- **ND2_hapNetwork** – ND2 haplotype ID in the Céspedes *et al* 2021 network.
- **kmN_wgri, kmN_HZ** – Northing distance (in km) from southernmost sample for whole-range and hybrid zone analyses.

**Table S2.** Optimization results for de novo assembly. Each tab corresponds to the three assemblies.

**Table S3.** Genetic diversity summary statistics by subspecies.
